# Magnesium Force Fields for OPC Water with Accurate Solvation, Ion-Binding, and Water-Exchange Properties: Successful Transfer from SPC/E

**DOI:** 10.1101/2021.12.07.471562

**Authors:** Kara K. Grotz, Nadine Schwierz

**Affiliations:** Department of Theoretical Biophysics, Max-Planck-Institute of Biophysics, Frankfurt am Main, Germany

## Abstract

Magnesium plays a vital role in a large variety of biological processes. To model such processes by molecular dynamics simulations, researchers rely on accurate force field parameters for Mg^2+^ and water. OPC is one of the most promising water models yielding an improved description of biomolecules in water. The aim of this work is to provide force field parameters for Mg^2+^ that lead to accurate simulation results in combination with OPC water. Using twelve different Mg^2+^ parameter sets, that were previously optimized with different water models, we systematically assess the transferability to OPC based on a large variety of experimental properties. The results show that the Mg^2+^ parameters for SPC/E are transferable to OPC and closely reproduce the experimental solvation free energy, radius of the first hydration shell, coordination number, activity derivative, and binding affinity toward the phosphate oxygens on RNA. Two optimal parameter sets are presented: *MicroMg* yields water exchange in OPC on the microsecond timescale in agreement with experiments. *NanoMg* yields accelerated exchange on the nanosecond timescale and facilitates the direct observation of ion binding events for enhanced sampling purposes.

## I. INTRODUCTION

Molecular dynamics simulations rely on accurate force field parameters for biomolecules, water molecules, and ions. It seems tempting to combine the force fields of the most promising water models with the most successful ion force fields in order to utilize the strengths of each parameter set. However, even for simple cations, the transferability of the ion parameters to different water models is limited. Different water models have a significant effect and can alter the thermodynamic and kinetic properties of the electrolyte solution considerably^1–3^. It is therefore crucial to assess whether the transfer of ion parameters to a different water model yields physically meaningful results. The aim of this work is to determine parameters for Mg^2+^ in OPC water that leverage the strengths of both force fields, reproduce a broad range of experimental properties, and lead to accurate simulation results of biomolecular systems.

Water constitutes the major part in simulations of membranes, proteins, and nucleic acids. Due to its distinct role, a large variety of water models exists which differ in their complexity, accuracy, and computational efficiency^4,5^. One of the most promising recent water models is the 4-site OPC model^5,6^. OPC water was developed to accurately reproduce the electrostatic properties of water. It is quoted to improve simulations of intrinsically disordered proteins^7,8^. Balancing the interactions between the water molecules and amino acids is particularly important for disordered proteins since most models tend to favor overly compact and collapsed structures^7,9^. Tian et al. recommend to employ OPC water in combination with their recently developed protein force field ff19SB^10^. In addition, the description of hydration of small molecules^6^ as well as the thermodynamics of ligand binding^11^ improves with OPC water. Moreover, small RNA fragments^12^ or central properties of DNA^13,14^ show significant improvement when simulated in OPC water.

The apparent success of OPC water in simulating biological system raises the immediate question which ion force field should be used to obtain reliable results. In particular, accurate parameters for Mg^2+^ are essential since these ions play a vital role in a large variety of physiological processes such as ATP hydrolysis^15^, cellular signaling^16,17^, or the catalytic activity of ribozymes^18–20^. Due to the prominent role of Mg^2+^, a variety of parameters for different water models exists today^1,21–34^. However, the development of accurate parameters for Mg^2+^ turned out to be notoriously difficult leading to various shortcomings of the parameters: (i) Simultaneously capturing the solvation free energy and the structure of the first hydration shell failed unless polarization effects were included explicitly^28,35^. (ii) Without further optimization, the existing parameters led to unrealistically slow exchange in the first hydration shell of Mg^2+^ rendering the simulation of ion binding events impossible^36^. (iii) The binding affinities to ion binding sites on biomolecules were typically overrated and needed further optimization^1,32,34,37^.

Recently, progress was made in tackling this problems by using a larger Lennard-Jones (LJ) parameter space and by modifying the standard combination rules^1,34^. In most biomolecular simulations, the pair potential between atoms *i* and *j* is modeled as sum of the Coulomb term and the 12-6 Lennard-Jones (LJ) potential,

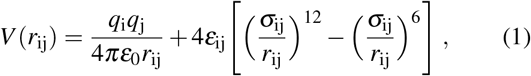

where *q*_i_ is the charge, *r*_ij_ the distance, and *ɛ*_0_ the dielectric constant of vacuum. The LJ term contains the interaction strength *ɛ*_ij_ and the diameter *σ*_ij_. Increasing the LJ parameter space renders the interaction between water and Mg^2+^ more attractive. This in turn allowed us to implicitly include polarization effects and to simultaneously reproduce the solvation free energy and the structure of the first hydration shell for 6 different water models^1,34^.

In addition, polarization effects provoke shortcomings of the standard combination rules for describing ion-ion interactions or the interaction of ions and biomolecules. By using scaling factors in the Lorentz-Berthelot combination rules^38–41^, the deviations from the standard combination rule can be taken into account. In particular, scaling parameters allowed us to reproduce activity derivatives over a broad range of MgCl_2_ concentrations and to correct the excessive binding of the Mg^2+^ ions to negatively charged groups on biomolecules^1,34^.

In the following, we systematically evaluate the transferability of those Mg^2+^ force field parameters, that were optimized previously in combination with different water models, to OPC water. Our results show that two parameter sets, called *microMg* and *nanoMg*, that were initially optimized in combination with SPC/E water, perform best in reproducing a broad variety of experimental properties in OPC water.

## II. METHODS

### A. Force field parameters

In our current work, we test the transferability of 12 different Mg^2+^ force field parameter sets^1,34^. Each parameter set was previously optimized in combination with the TIP3P^42^, SPC/E^43^, TIP3P-fb^44^, TIP4P/2005^45^, TIP4P-Ew^46^, and TIP4P-D^9^ water model. In the following, we refer to these parameter sets as TIP3P-, TIP3P-fb-, TIP4P/2005-, TIP4P-Ew-, or TIP4P-D-optimized parameter sets. The corresponding Mg^2+^ force field parameters are available from github https://github.com/bio-phys/Magnesium-FFs and https://github.com/bio-phys/optimizedMgFFs or from Table S2 in the SI. Note that for each water model two parameter sets exist, labeled *microMg* and *nanoMg*. The *microMg* parameter set reproduces the experimental water exchange rate on the microsecond time scale while the *nanoMg* parameter set yields accelerated exchanges on the nanosecond timescale. The parameters of the water models are listed in Table S1 of the SI.

The results upon transferring those force field parameters to OPC are compared to the Mg^2+^ force field parameters by Li and coworkers^33^ that were recently optimized in combination with OPC water. In the following, we refer to them as Li-Merz OPC-optimized parameters. In the work by Li et al.^33^, two different parameter sets are given. The first set of parameters is based on the commonly used 12-6 Lennard-Jones interaction potential. The second set of parameters is based on the so-called 12-6-4 potential which introduces an additional term to mimic polarization effects^28^.

### B. Combination rules

To describe the Mg^2+^-water interactions, we use unmodified Lorentz-Berthelot combination rules

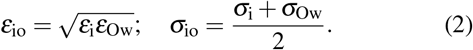

Note that the parameters *ɛ*_Ow_ and *σ*_Ow_ correspond to the ones for OPC water in this current work (see Table S1). The values for *ɛ*_io_ and *σ*_io_ therefore deviate from the values with the respective original water models^1,34^ while the values for *ɛ*_ii_ and *σ*_ii_ remain unchanged (Table I).

**TABLE I.** Optimal force field parameters for simulations of Mg^2+^ in OPC water. The parameters were previously optimized for SPC/E^1^ and reproduce a broad range of experimental properties when transferred to OPC (Table II). *σ*_ii_, *ɛ*_ii_, *σ*_io_, *ɛ*_io_ are the ion-ion and ion-water LJ parameters. 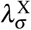 and 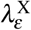 are the scaling factors for the Lorentz-Berthelot combination rules (eq 3) for the interaction with Cl or the RNA atoms. Note that the scaling factors are only valid in combination with the Cl^−^ parameters from Smith-Dang^49^ for SPC/E water, and the parmBSC0*χ*_OL3_ RNA parameters^55–57^.

|  |  | *microMg* | *nanoMg* |
| --- | --- | --- | --- |
| $\sigma_{ii}$ | [nm] | 0.1036 | 0.1046 |
| $\epsilon_{ii}$ | [kJ/mol] | 290.58 | 470.70 |
| $\sigma_{io}$ | [nm] | 0.2101 | 0.2106 |
| $\epsilon_{io}$ | [kJ/mol] | 16.08 | 20.47 |
| $\lambda_{\sigma}^{\text{Cl}}$ | | 1.59 | 1.59 |
| $\lambda_{\epsilon}^{\text{Cl}}$ | | 0.10 | 0.10 |
| $\sigma_{\text{MgCl}}$ | [nm] | 0.4316 | 0.4325 |
| $\epsilon_{\text{MgCl}}$ | [kJ/mol] | 1.0989 | 1.3986 |
| $\lambda_{\sigma}^{\text{RNA}}$ | | 1.1019 | 1.1107 |
| $\lambda_{\epsilon}^{\text{RNA}}$ | | 0.4856 | 0.3300 |
| $\sigma_{\text{MgOP}}$ | [nm] | 0.2202 | 0.2225 |
| $\epsilon_{\text{MgOP}}$ | [kJ/mol] | 7.7589 | 6.7111 |

**TABLE II.**
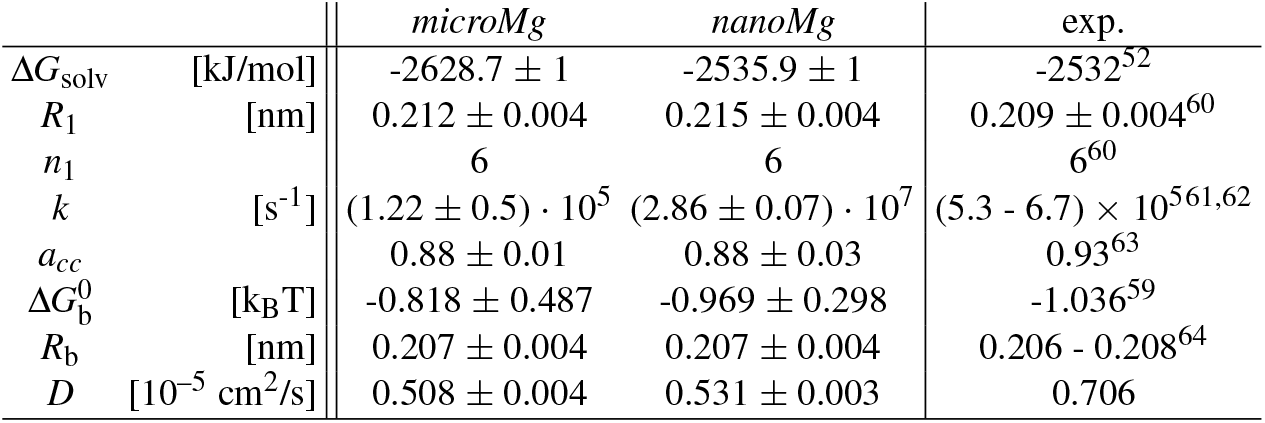
Results for single-ion, ion-ion and ion-RNA properties in OPC for the optimal Mg^2+^ parameters (Table I). Solvation free energy of neutral MgCl_2_ ion pairs Δ*G*_solv_, Mg^2+^-oxygen distance in the first hydration shell *R*_1_, coordination number of the first hydration shell *n*_1_, water exchange rate from the first hydration shell *k*, binding affinity toward the phosphate oxygen of DMP 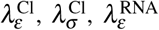, Mg^2+^-phosphate oxygen distance in inner-sphere coordination *R*_b_, and *a_cc_* the activity derivative of a MgCl_2_ solution at 0.25 M concentration. The experimental value for 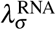 is derived from the log stability constant (log *K* = 0.45) given in ref.^59^. *D* is the self-diffusion coefficient.

To model the Mg^2+^-Cl^−^ and Mg^2+^-RNA interactions, we introduce adjustable scaling parameters 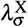 and 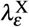 in the Lorentz-Berthelot combination, similar to previous works^1,32,34,38–41^. With this, the Lorentz-Berthelot combination rules have the following form

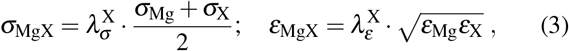

where X represents the Cl^−^ anion or the atoms of an RNA. The scaling parameters allow us to take some of the effects of polarizablity into account which can cause deviations from the standard combination rule. At the same time, the Mg^2+^-water interaction remains unchanged such that the solvation free energy, the structural properties of the first hydration shell, and the rate of water exchange are not affected.

### C. Simulation setup

All simulations were done with GROMACS^47^ (version 2020) with the exception of the 12-6-4 interaction potentials since this interaction form is not readily available in GROMACS. Therefore, simulations with the 12-6-4 interaction potential were done with AMBER^48^ (version 2018). We used different simulation setups to calculate the different physical properties. The details of the different setups are described in Section S3 of the SI.

### D. Transferability of Mg^2+^ parameters to OPC water

The simulations to evaluate the transferability of different force field parameters to OPC water were done in three consecutive steps.

#### 1. Solvation free energy, radius, and coordination number

In the first step, we calculated all single-ion properties for the twelve transferred and for the two Li-Merz parameter sets. In order to provide neutral systems, we used Cl^−^ as counter ion. In particular, we used the Cl^−^ parameters by Mamatkulov-Schwierz^30^ for TIP3P, Smith-Dang^49^ for SPC/E, and Grotz-Schwierz^1^ for TIP3P-fb, TIP4P/2005, TIP4P-Ew, and TIP4P-D.

We calculated the solvation free energy of neutral MgCl_2_ ion pairs which includes the enthalpic and entropic contribution of the ions in water. In the computation, we took correction terms for finite size^50^ and pressure^51^ into account. Note that the term for interfacial crossing^51^ cancels for neutral ion pairs. For the OPC-optimized 12-6 and 12-6-4 parameters, the experimental value for Cl^-^^52^ was added to the reported literature values^33^ to obtain a neutral ion pair. Further details on the calculation of the solvation free energy can be found in Section S4 of the SI.

In addition, we considered the radius and coordination number as the most important structural parameters to characterize the first hydration shell. In the computation of radius and coordination number, the Sengupta-Merz 12-6 and 12-6-4 parameters^53^, that were optimized in combination with OPC, were considered as corresponding Cl^−^ parameters for the Li-Merz Mg^2+^ parameters.

#### 2. Water exchange rate

In the second step, we selected the 8 parameter sets that performed best in the previous step and computed the water exchange rates in the first hydration shell. The exchange rates were obtained by counting transitions of water molecules between the first and second hydration shell from long trajectories in MgCl_2_. Note, however, that for some force field parameters water exchange becomes so rare that the rate could not be calculated from straight forward simulations^36^. While this limitation did not affect the transferred parameters, exchanges with the 12-6 and 12-6-4 Li-Merz parameters were rare. In both cases, water exchange was observed to be unrealistically slow compared to the experimental results and the rate could not be determined with good statistics (12-6) or not at all (12-6-4). More details on these computations can be found in Section S5 of the SI.

#### 3. Activity derivative and binding affinity

In the last step, the activity derivatives in MgCl_2_ and the binding affinity toward one of the non-bridging phosphate oxygens of RNA were calculated. Note that in both cases the respective modified Lorentz-Berthelot combination rules (eq 3) need to be used for accurate results. The corresponding scaling factors, 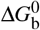, and 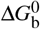 are given in Tables I and S2. The activity derivatives were obtained from straight forward simulations in MgCl_2_ solutions with different concentrations and Kirkwood-Buff theory (see Section S6 in SI for more details).

To calculate the binding affinity of Mg^2+^, we used the dimethylphosphate (DMP) as a simple model system containing the two non-bridging phosphate oxygen atoms. The GAFF^54^ and AMBER RNA force field parameters (parmBSC0*χ*_OL3_^55–57^) were used. Further details on the force field parameters for DMP can be found in ref.^34^. For the final two parameter sets the binding affinity was obtained from two different methods, namely by integrating potentials of mean force and via alchemical transformations. More details on the calculations of the binding affinities can be found in Section S7 of the SI.

### E. Free energy profiles

To gain further insight into water exchange and Mg^2+^ binding to DMP, we calculated the one-dimensional free energy profiles. For water exchange, we used umbrella sampling to obtain the free energy profiles as a function of the Mg^2+^-water oxygen distance. For Mg^2+^ binding, we used umbrella sampling to obtain the free energy profiles as a function of the Mg^2+^-phosphate oxygen distance. Further details can be found in Section S5 and S7.1 of the SI. The barrier heights are listed in Table S6 for water exchange and in Table S7 for Mg^2+^ binding to DMP.

### F. Diffusion coefficient

For the final parameter sets, the self-diffusion coefficient was calculated from an additional 10 ns NVT simulation of the single Mg^2+^ ion in OPC water. The first nanosecond was excluded from the analysis for equilibration. The self-diffusion coefficient was calculated from the slope of the mean-square displacement. After a brief initial period, the mean-square displacement grows linearly and the diffusion coefficient is estimated from a straight line fit. The diffusion coefficient was corrected for system size effects^58^,

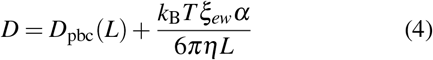

where *L* is the box length, *D*_pbc_ the computed self-diffusion coefficient, *D* the diffusion coefficient for the infinite non-periodic system, *k_B_* the Boltzmann constant, *T* the absolute temperature, and *ξ_ew_* = 2.837297 the self-term for a cubic lattice. The empirical parameter *α* was set to 1.0. Since the solvent viscosity *η* for OPC water has not been reported in the literature, we used the experimental value *η* = 8.91 · 10^−4^ kg m^−1^s^−14^.

## III. RESULTS

The aim of this work is to evaluate the transferability of different Mg^2+^ force fields to OPC water. In particular, we calculated all physical properties that were targeted in the initial optimization including the solvation free energy, the distance to water oxygens in the first hydration shell, the hydration number, the activity coefficient derivative in MgCl_2_ solutions, and the binding affinity and distance to the non-bridging phosphate oxygens on nucleic acids. Finally, we selected two parameter sets (Table I), *microMg* and *nanoMg*, that reproduce the broad range of experimental properties (Table II) and lead to accurate simulation results in OPC water and are recommended for conventional or enhanced sampling purposes, respectively.

### A. Solvation free energy, Mg^2+^-water distance, and coordination number

To evaluate whether the Mg^2+^ parameters, that were optimized in combination with different 3- and 4-site water models, are transferable, we calculated the solvation free energy of MgCl_2_ ion pairs in OPC water. Note that the usage of neutral ion pairs is more robust since it does not rely on conflicting experimental results for the hydration free energy of a proton^52,65,66^. Figure 1A shows the deviation of the calculated solvation free energy from the experimental results ΔΔ*G*_solv_ for the transferred parameters. For comparison, the values for the OPC-optimized 12-6 and 12-6-4 Li-Merz parameters^33^ are shown. The simulations with the parameters transferred from SPC/E to OPC water yield the smallest deviation from experiments and perform even slightly better than those optimized directly within OPC.

**FIG. 1.**
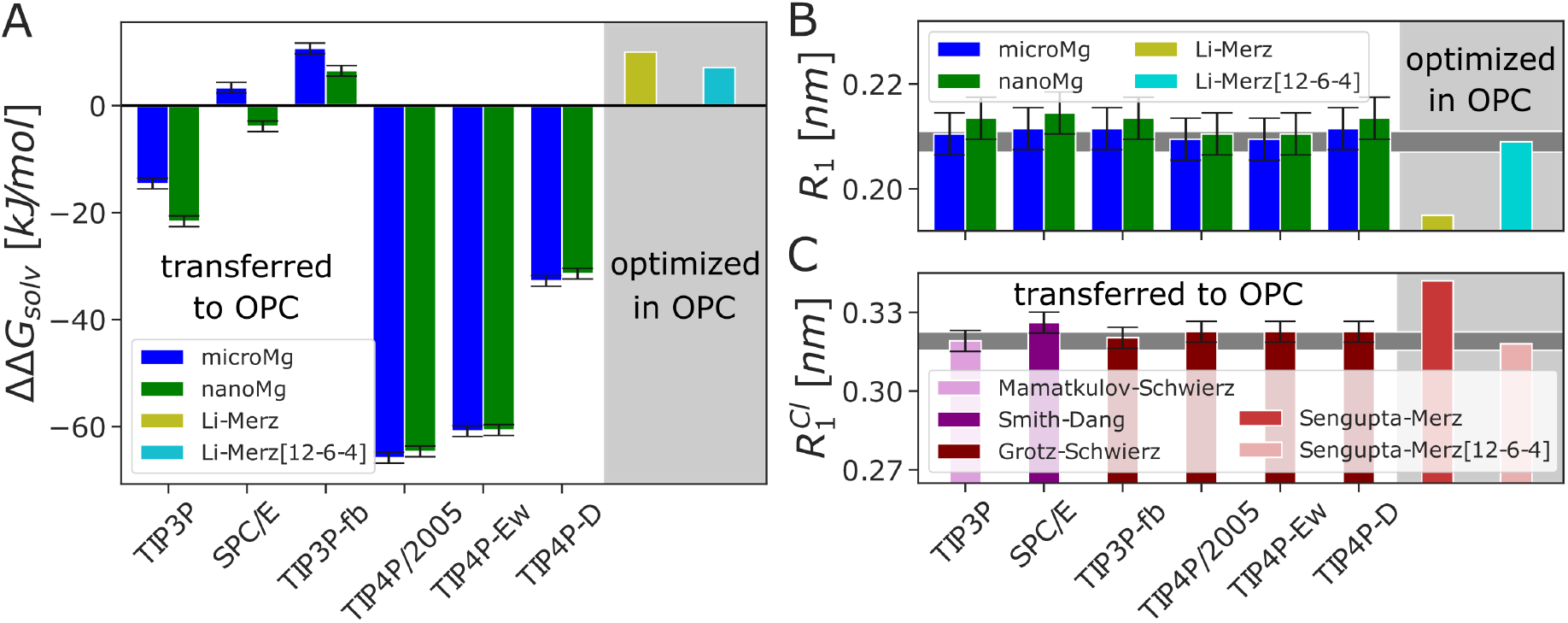
(A) Deviation of the solvation free energy from the experimental result^52^ ΔΔ*G*_solv_ for neutral MgCl_2_ ion pairs. The results are obtained by transferring the Mg^2+^ parameters obtained in the different water models (TIP3P, SPC/E, TIP3P-fb, TIP4P/2005, TIP4P-Ew, or TIP4P-D) to OPC. For comparison, the OPC-optimized 12-6 and 12-6-4 Li-Merz^33^ parameters are shown. Since ref.^33^ only contains values for Mg^2+^, the experimental value for Cl^−^^52^ was added to obtain a neutral ion pair. (B) Mg^2+^ - oxygen distance *R*_1_ in the first hydration shell. (C) Cl^−^ - oxygen distance 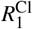 of the first hydration shell obtained with different force fields from the literature^1,30,49,53^. The gray horizontal bar in (B,C) corresponds to the experimental value^60^.

Interestingly, ΔΔ*G*_solv_ is smaller for the parameters transferred from 3-site waters (SPC/E, followed by TIP3P-fb and TIP3P). 4-site water models (TIP4P-D, followed by TIP4P-Ew, and TIP4P/2005) lead to larger deviations despite the fact that OPC is also a 4-site model (Table S4). This result has not been expected based on previous work which showed reasonable transferability of ion parameters within water models of the same complexity^1,2,67^.

The size of the first hydration shell, measured by the distance between Mg^2+^ and the water oxygens *R*_1_, agrees well with the experimental results for all parameter sets transferred to OPC water (Figure 1B, Table S4). On the other hand, the OPC-optimized 12-6 model by Li-Merz significantly underestimates *R*_1_. This shortcoming led to the development of the 12-6-4 model^28^. Not surprisingly, the OPC-optimized 12-6-4 model^33^ provides much better agreement (Figure 1B).

For all parameter sets, the coordination number in OPC water is 6 and precisely matches the experimental result (Table S4).

Similarly, the simulations with Cl^−^ parameter from different water models transferred to OPC closely reproduce *R*_1_ (Figure 1C, Table S5). On the other hand, the OPC-optimized 12-6 Sengupta-Merz parameters deviate noticeably from the experimental value while the 12-6-4 version yields good agreement.

In summary, the six Mg^2+^ parameter sets that were previously optimized for the 3-site water models (SPC/E, followed by TIP3P-fb and TIP3P) and their corresponding three Cl^−^ parameter sets yield the best results for the solvation free energy and structure of the first hydration shell when transferred to OPC water and are used in the subsequent steps.

### B. Water exchange rate

Including water exchange rates in the evaluation of the transferability is important to correctly capture ion binding and to avoid shortcomings due to unrealistically slow exchange dynamics as observed previously^36^. In particular, we evaluated the capability of *microMg* in reproducing the experimental exchange rate and the capability of *nanoMg* to speed up the association or dissociation processes for application in enhanced sampling simulations. Figure 2 shows the water exchange rate for the six parameter sets transferred to OPC water in comparison to the OPC-optimized Li-Merz parameters and the experimental results^61,62^. All transferred parameter sets yield a lower exchange rate in OPC water compared to the original water reflecting the influence of the water model on the exchange kinetics^3^. Still, the TIP3P-fb and SPC/E-optimized *microMg* parameters yield the same order of magnitude as in experiments and are therefore considered reasonably accurate (Table S6). In addition, the transferred *microMg* parameters perform significantly better compared to the 12-6 and 12-6-4 Li-Merz parameters. Note, however, that there is a large uncertainty in the rate for the 12-6 Li-Merz parameters as the exchange dynamics is very slow with only 12 exchanges in 2 *μ*s. With the 12-6-4 parameters, the exchange is even slower and not a single exchange event could be observed (Table S6). Even though it is surprising that the 12-6-4 parameters lead to a slower exchange compared to the 12-6 parameters despite including polarization effects via the *r*^−4^ term, the behavior is well-reflected in the one-dimensional free energy profiles (Figure S1D).

**FIG. 2.**
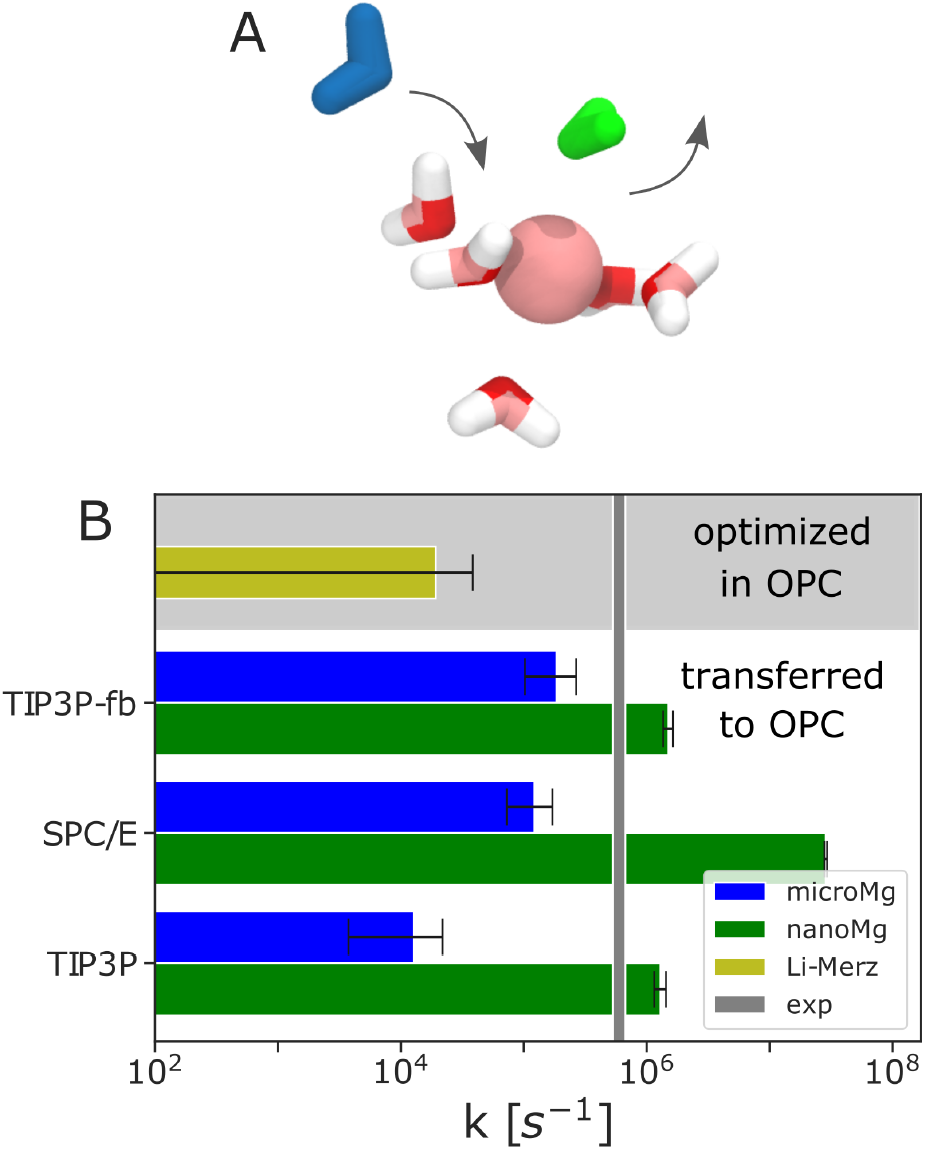
(A) Snapshot of a Mg^2+^ ion including the first hydration shell during a direct water exchange: The incoming water is shown in blue, the outgoing water in green. (B) Water exchange rates calculated from 2 *μ*s of 1 M MgCl_2_ trajectories for the different force field parameters. The gray vertical bar indicates the experimental rate^61,62^. Errors are calculated from block averaging by dividing the trajectory into 2 blocks.

For the *nanoMg* parameter sets, significant differences are observed: With the SPC/E-optimized parameters in OPC, exchanges happen on the order of 10^7^/s and are therefore almost as fast as in SPC/E (10^8^/s)^1^. With the TIP3P-fb-optimized and TIP3P-optimized *nanoMg* parameters transferred to OPC, the maximum rate is on the same order of magnitude (10^6^/s) and two orders of magnitude slower (10^8^/s to 10^6^/s), respectively. The SPC/E-optimized *nanoMg* parameters in OPC are therefore most beneficial for enhanced sampling of ion binding in OPC water.

In summary, based on the water exchange rate, the SPC/E- and TIP3P-fb-optimized parameters yield the best agreement with experimental results (*microMg* sets) while the SPC/E-optimized *nanoMg* parameters yield the highest acceleration of exchanges.

### C. Activity derivative and binding affinity to RNA

Finally, we evaluate the transferability based on the activity derivative and the binding affinity and distance to the phosphate oxygens on nucleic acids. The activity derivative *a*_cc_ is important in the evaluation of the transferability since it gives information on the balance between ion-ion and ion-water interactions^38^. Figure 3A shows that the TIP3P-, SPC/E- and TIP3P-fb-optimized parameters reproduce *a*_cc_ over a broad concentration range in OPC water.

**FIG. 3.**
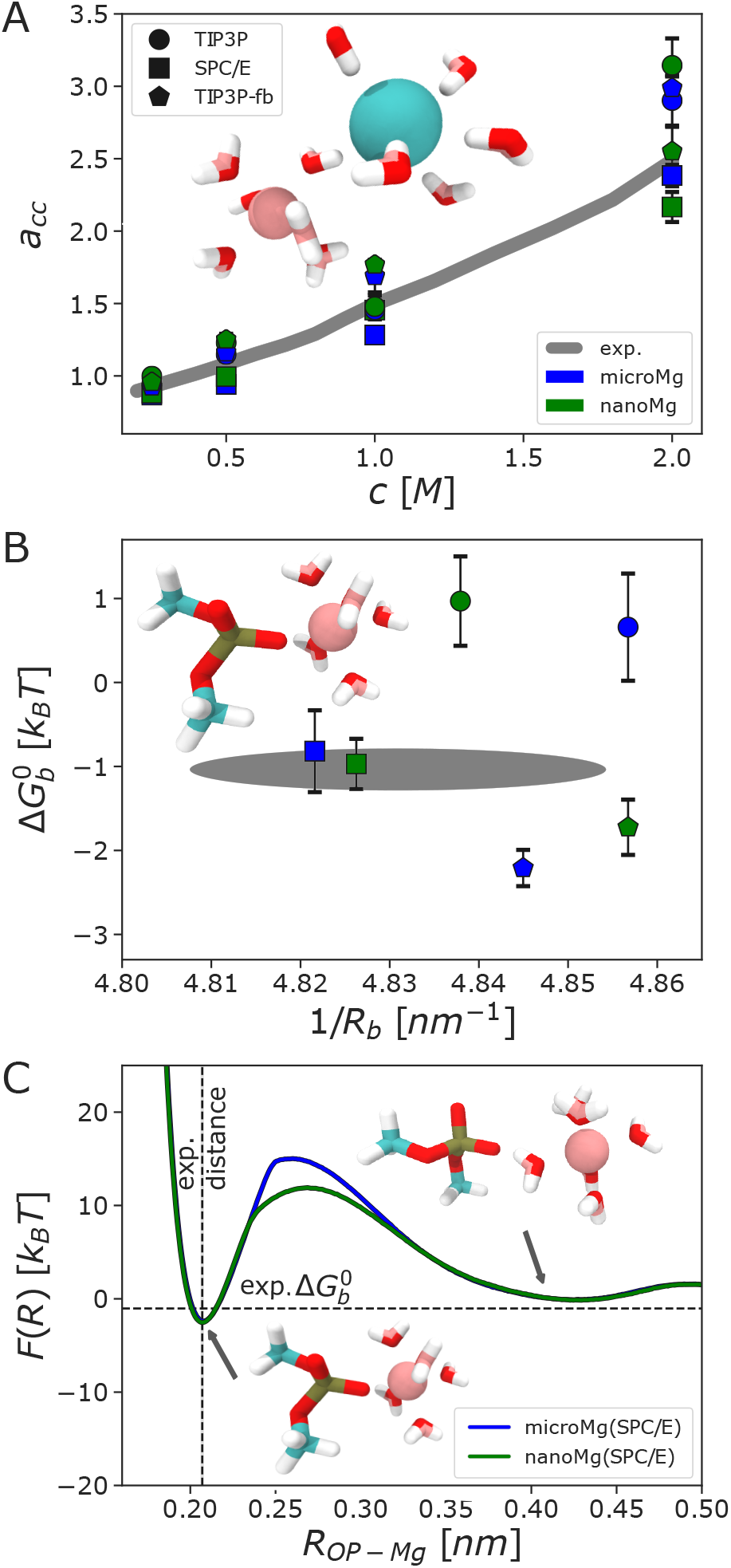
(A) Activity derivative *a*_cc_ as function of the MgCl_2_ concentration for selected force field parameters. The inset shows a Mg^2+^-Cl^−^ pair and its first hydration shells. The gray line corresponds to the experimental value from ref.^63^. (B) Binding affinity 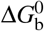 in correlation with the inverse of the binding distance 1/*R*_b_ toward the phosphate oxygen. The inset shows a simulation snapshot of the DMP molecule with one Mg^2+^ ion in inner-sphere coordination. The gray area represents the experimental values from refs.^59,64^. (C) Free energy profiles *F*(*R*) along the Mg^2+^-phosphate oxygen distance for the optimal parameters sets in OPC water. The insets show simulation snapshots of Mg^2+^ in the two stable states.

The binding affinity 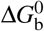 and distance *R*_b_ provide insight into the accuracy of the Mg^2+^-RNA interactions and are shown in Figure 3B. The SPC/E-optimized parameters precisely match 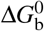 and *R*_b_ in OPC water, whereas parameters for TIP3P overestimate and those for TIP3P-fb underestimate 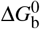.

In summary, the SPC/E-optimized *microMg* and *nanoMg* parameters yield the best agreement with the experimental results in OPC water.

Figure 3C provides additional insight into the ion binding process from the free energy profiles along the Mg^2+^-phosphate oxygen distance for the SPC/E-optimized parameters in OPC. The two minima correspond to the inner- and outer-sphere coordination of Mg^2+^ and are identical for *microMg* and *nanoMg*, as expected. The energetic barrier, on the other hand, is significantly lower for *nanoMg* reflecting its enhanced ion binding kinetics.

## IV. CONCLUSION

OPC water has proven to be one of the most promising 4-site water models^6^ due to its improved description of small molecules^6,11^, proteins^7,8,10^, and nucleic acids^12–14^. In this work, we evaluate the transferability of twelve different Mg^2+^ force fields to OPC water based on solvation free energy, size of the first hydration shell, hydration number, water exchange rate, activity derivative, and binding affinity to the phosphate oxygens on nucleic acids. Our results show that the force field parameters, that were previously optimized with SPC/E^1^, are best suited to reproduce the broad range of solution properties in OPC water. In addition, the results show that the transferred parameters are compatible or better than OPC-optimized 12-6 and 12-6-4 parameters^33^ in reproducing the experimental solvation free energy and water exchange rate.

Moreover, the SPC/E-optimized parameters closely reproduce the binding affinity of Mg^2+^ toward the phosphate oxygen of RNA. Matching the binding affinity of metal cations to specific ion binding sites is particularly important since it improves the agreement between experiments and simulations for the structure and properties of larger nucleic acid systems^68^.

In summary, the SPC/E-optimized parameter sets *microMg* and *nanoMg* provide an efficient and highly accurate model for the simulation of Mg^2+^ in OPC water.

We recommend to use *microMg* to obtain accurate water exchange kinetics comparable to experimental results on the microsecond timescale. Further, we recommend to use the *nanoMg* parameters to yield accelerated exchange kinetics and ion-binding in enhanced sampling setups.

## V. ACKNOWLEDGEMENTS

We acknowledge financial support from the DFG (Emmy Noether program, Grant No. 315221747 and the CRC902). GOETHE HLR is acknowledged for supercomputing access.

## VI. SUPPORTING INFORMATION

Further details of the simulations and the analysis are provided in the supplementary material: Simulation setup, parameters of different water models and Mg^2+^ force fields, details on the computation of the solvation free energy, water exchange rates, activity derivatives, radial distribution functions between Mg^2^ and Cl^2^ for the different models, and binding affinities. The Mg^2+^ force field parameters are also available at https://github.com/bio-phys/MgFF_OPC.

## VII. DATA AVAILABILITY

The data that support the finding of this study are available from the corresponding author upon reasonable request.

## Notes

https://github.com/bio-phys/MgFF_OPC

### Competing Interest Statement

The authors have declared no competing interest.

## References

1 K. K. Grotz and N. Schwierz, “Optimized Magnesium Force Field Parameters for Biomolecular Simulations with Accurate Solvation, Ion-Binding, and Water-Exchange Properties in SPC/E, TIP3P-fb, TIP4P/2005, TIP4P-Ew, and TIP4P-D,” J. Chem. Theory Comput. 19, 526–537 (2022).

2 M. F. Döpke, O. A. Moultos, and R. Hartkamp, “On the transferability of ion parameters to the TIP4P/2005 water model using molecular dynamics simulations,” J. Chem. Phys. 152, 024501 (2020).

3 S. Falkner and N. Schwierz, “Kinetic pathways of water exchange in the first hydration shell of magnesium: Influence of water model and ionic force field,” J Chem. Phys. 155, 084503 (2021).

4 C. Vega and J. L. F. Abascal, “Simulating water with rigid non-polarizable models: a general perspective,” Phys. Chem. Chem. Phys. 13, 19663–19688 (2011).

5 S. P. K. Pathirannahalage, N. Meftahi, A. Elbourne, A. C. G. Weiss, C. F. McConville, A. Padua, D. A. Winkler, M. Costa Gomes, T. L. Greaves, Q. A. Besford, T. C. Le, and A. J. Christofferson, “A systematic comparison of the structural and dynamic properties of commonly used water models for molecular dynamics simulations,” J. Chem. Inf. Model 61, 4521–4536 (2021).

6 S. Izadi, R. Anandakrishnan, and A. V. Onufriev, “Building water models: A different approach,” J. Phys. Chem. Lett. 5, 3863–3871 (2014), arXiv:1408.1679.

7 P. S. Shabane, S. Izadi, and A. V. Onufriev, “General Purpose Water Model Can Improve Atomistic Simulations of Intrinsically Disordered Proteins,” J. Chem. Theory Comput. 15, 2620–2634 (2019).

8 D. Pantoja-Uceda, J. L. Neira, L. M. Contreras, C. A. Manton, D. R. Welch, and B. Rizzuti, “The isolated C-terminal nuclear localization sequence of the breast cancer metastasis suppressor 1 is disordered,” Arch. Biochem. Biophys. 664, 95–101 (2019).

9 S. Piana, A. G. Donchev, P. Robustelli, and D. E. Shaw, “Water dispersion interactions strongly influence simulated structural properties of disordered protein states,” J. Phys. Chem. B 119, 5113–5123 (2015).

10 C. Tian, K. Kasavajhala, K. A. Belfon, L. Raguette, H. Huang, A. N. Migues, J. Bickel, Y. Wang, J. Pincay, Q. Wu, and C. Simmerling, “FF19SB: Amino-Acid-Specific Protein Backbone Parameters Trained against Quantum Mechanics Energy Surfaces in Solution,” J. Chem. Theory Comput. 16, 528–552 (2020).

11 K. Gao, J. Yin, N. M. Henriksen, A. T. Fenley, and M. K. Gilson, “Binding Enthalpy Calculations for a Neutral Host-Guest Pair Yield Widely Divergent Salt Effects across Water Models,” J. Chem. Theory Comput. 11, 4555–4564 (2015).

12 C. Bergonzo and T. E. Cheatham, “Improved Force Field Parameters Lead to a Better Description of RNA Structure,” J. Chem. Theory Comput. 11, 3969–3972 (2015).

13 F. Häse and M. Zacharias, “Free energy analysis and mechanism of base pair stacking in nicked DNA,” Nucleic Acids Res. 44, 7100–7108 (2016).

14 R. Galindo-Murillo, J. C. Robertson, M. Zgarbová, J. Šponer, M. Otyepka, P. Jurečka, and T. E. Cheatham, “Assessing the Current State of Amber Force Field Modifications for DNA,” J. Chem. Theory Comput. 12, 4114–4127 (2016).

15 N. H. Williams, “Magnesium Ion Catalyzed ATP Hydrolysis,” J. Am. Chem. Soc. 122, 12023–12024 (2000).

16 J. H. de Baaij, J. G. Hoenderop, and R. J. Bindels, “Magnesium in man: Implications for health and disease,” Physiol. Rev. 95, 1–46 (2015).

17 A. Stangherlin and J. S. O’Neill, “Signal Transduction: Magnesium Manifests as a Second Messenger,” Curr. Biology 28, R1403–R1405 (2018).

18 J. A. Cowan, “Structural and catalytic chemistry of magnesium-dependent enzymes,” Biometals 15, 225–235 (2002).

19 A. Pyle, “Metal ions in the structure and function of RNA,” J. Biol. Inorg. Chem. 7, 679–690 (2002).

20 R. K. O. Sigel and A. M. Pyle, “Alternative Roles for Metal Ions in Enzyme Catalysis and the Implications for Ribozyme Chemistry,” Chem. Rev. 107, 97–113 (2007).

21 J. Åqvist, “Ion-water interaction potentials derived from free energy perturbation simulations,” J. Phys. Chem. 94, 8021–8024 (1990).

22 C. S. Babu and C. Lim, “Empirical force fields for biologically active divalent metal cations in water,” J. Phys. Chem. A 110, 691–699 (2006).

23 D. M. Mayaan, Evelyn; Moser, Adam; MacKerell, Aalexander D. Jr., York, “CHARMM Force Field Parameters for Simulation of Reactive Intermediates in Native and Thio-Substituted Ribozymes,” J. Comput. Chem. 28, 495–507 (2007).

24 E. Duboué-Dijon, P. Delcroix, H. Martinez-Seara, J. Hladílková, P. Coufal, T. Křížek, and P. Jungwirth, “Binding of Divalent Cations to Insulin: Capillary Electrophoresis and Molecular Simulations,” J. Phys. Chem. B 122, 5640–5648 (2018).

25 O. Allnér, L. Nilsson, and A. Villa, “Magnesium Ion-Water Coordination and Exchange in Biomolecular Simulations,” J. Chem. Theory Comput. 8, 1493–1502 (2012).

26 S. Mamatkulov, M. Fyta, and R. R. Netz, “Force fields for divalent cations based on single-ion and ion-pair properties,” J. Chem. Phys. 138, 24505 (2013).

27 P. Li, B. P. Roberts, D. K. Chakravorty, and K. M. Merz, “Rational Design of Particle Mesh Ewald Compatible Lennard-Jones Parameters for +2 Metal Cations in Explicit Solvent,” J. Chem. Theory Comput. 9, 2733–2748 (2013).

28 P. Li and K. M. Merz, “Taking into Account the Ion-Induced Dipole Interaction in the Nonbonded Model of Ions,” J. Chem. Theory Comput. 10, 289–297 (2014).

29 M. T. Panteva, G. M. Giambasu, and D. M. York, “Comparison of structural, thermodynamic, kinetic and mass transport properties of Mg^2+^ ion models commonly used in biomolecular simulations,” J. Comput. Chem. 36, 970–982 (2015).

30 S. Mamatkulov and N. Schwierz, “Force fields for monovalent and divalent metal cations in TIP3P water based on thermodynamic and kinetic properties,” J. Chem. Phys. 148, 74504 (2018).

31 H. T. Nguyen, N. Hori, and D. Thirumalai, “Theory and simulations for RNA folding in mixtures of monovalent and divalent cations,” Proc. Natl. Acad. Sci. 116, 21022–21030 (2019).

32 S. Cruz-León, K. K. Grotz, and N. Schwierz, “Extended magnesium and calcium force field parameters for accurate ion-nucleic acid interactions in biomolecular simulations,” J. Chem. Phys. 154, 171102 (2021).

33 Z. Li, L. F. Song, P. Li, and K. M. Merz, “Systematic Parametrization of Divalent Metal Ions for the OPC3, OPC, TIP3P-FB, and TIP4P-FB Water Models,” J. Chem. Theory Comput. 16, 4429–4442 (2020).

34 K. K. Grotz, S. Cruz-León, and N. Schwierz, “Optimized Magnesium Force Field Parameters for Biomolecular Simulations with Accurate Solvation, Ion-Binding, and Water-Exchange Properties,” J. Chem. Theory Comput. 17, 2530–2540 (2021).

35 I. M. Zeron, J. L. Abascal, and C. Vega, “A force field of Li^+^, Na^+^, K^+^, Mg^2+^, Ca^2+^, Cl^−^, and SO_4_^2−^ In aqueous solution based on the TIP4P/2005 water model a,” J. Chem. Phys. 151, 134504 (2019).

36 N. Schwierz, “Kinetic pathways of water exchange in the first hydration shell of magnesium,” J. Chem. Phys. 152, 224106 (2020).

37 M. T. Panteva, G. M. Giambacsu, and D. M. York, “Force field for Mg^2+^, Mn^2+^, Zn^2+^, and Cd^2+^ ions that have balanced interactions with nucleic acids,” J. Phys. Chem. B 119, 15460–15470 (2015).

38 M. B. Gee, N. R. Cox, Y. Jiao, N. Bentenitis, S. Weerasinghe, and P. E. Smith, “A Kirkwood-Buff Derived Force Field for Aqueous Alkali Halides,” J. Chem. Theory Comput. 7, 1369–1380 (2011).

39 M. Fyta and R. R. Netz, “Ionic force field optimization based on single-ion and ion-pair solvation properties: Going beyond standard mixing rules,” J. Chem. Phys. 136, 124103 (2012).

40 S. Weerasinghe and P. E. Smith, “A Kirkwood-Buff derived force field for sodium chloride in water,” J. Chem. Phys. 119, 11342–11349 (2003).

41 B. Hess and N. F. A. van der Vegt, “Cation specific binding with protein surface charges,” Proc. Natl. Acad. Sci. 106, 13296–13300 (2009).

42 W. L. Jorgensen, J. Chandrasekhar, J. D. Madura, R. W. Impey, and M. L. Klein, “Comparison of simple potential functions for simulating liquid water,” J. Chem. Phys. 79, 926–935 (1983).

43 H. J. Berendsen, J. R. Grigera, and T. P. Straatsma, “The missing term in effective pair potentials,” J. Phys. Chem. 91, 6269–6271 (1987).

44 L. P. Wang, T. J. Martinez, and V. S. Pande, “Building force fields: An automatic, systematic, and reproducible approach,” J. Phys. Chem. Lett. 5, 1885–1891 (2014).

45 J. L. Abascal and C. Vega, “A general purpose model for the condensed phases of water: TIP4P/2005,” J. Chem. Phys. 123, 234505 (2005).

46 H. W. Horn, W. C. Swope, J. W. Pitera, J. D. Madura, T. J. Dick, G. L. Hura, and T. Head-Gordon, “Development of an improved four-site water model for biomolecular simulations: TIP4P-Ew,” J. Chem. Phys. 120, 9665–9678 (2004).

47 M. J. Abraham, T. Murtola, R. Schulz, S. Páll, J. C. Smith, B. Hess, and E. Lindah, “Gromacs: High performance molecular simulations through multi-level parallelism from laptops to supercomputers,” SoftwareX 1-2, 19–25 (2015).

48 D. A. Case, K. Belfon, I. Y. Ben-Shalom, S. R. Brozell, D. S. Cerutti, T. E. I. Cheatham, V. W. D. Cruzeiro, T. Darden, R. E. Duke, G. Giambasu, M. K. Gilson, H. Gohlke, A. W. Goetz, R. Harris, P. A. Izadi, S. Izmailov, K. Kasavajhala, A. Kovalenko, R. Krasny, T. Kurtzman, T. S. Lee, S. LeGrand, P. Li, C. Lin, J. Liu, T. Luchko, R. Luo, V. Man, K. M. Merz, Y. Miao, O. Mikhailovskii, G. Monard, H. Nguyen, A. Onufriev, F. Pan, S. Pantano, R. Qi, D. R. Roe, A. Roitberg, C. Sagui, S. Schott-Verdugo, J. Shen, C. L. Simmerling, N. R. Skrynnikov, J. Smith, J. Swails, R. C. Walker, J. Wang, L. Wilson, R. M. Wolf, X. Wu, Y. Xiong, Y. Xue, D. M. York, and P. A. Kollman, “Amber 2018,” (2018).

49 D. E. Smith and L. X. Dang, “Computer simulations of NaCl association in polarizable water,” J. Chem. Phys. 100, 3757–3766 (1994).

50 G. Hummer, L. R. Pratt, and A. E. García, “Free Energy of Ionic Hydration,” J. Phys. Chem. 100, 1206–1215 (1996).

51 D. Horinek, S. I. Mamatkulov, and R. R. Netz, “Rational design of ion force fields based on thermodynamic solvation properties,” J. Chem. Phys. 130, 124507 (2009).

52 Y. Marcus, Ion Properties (Marcel Dekker, Inc., New York, Basel, 1997).

53 A. Sengupta, Z. Li, L. F. Song, P. Li, and K. M. Merz, “Parameterization of Monovalent Ions for the OPC3, OPC, TIP3P-FB, and TIP4P-FB Water Models,” J. Chem. Inf. Model 61, 869–880 (2021).

54 J. Wang, R. M. Wolf, J. W. Caldwell, P. A. Kollman, and D. A. Case, “Development and testing of a general Amber force field,” J. Comput. Chem. 25, 1157–1174 (2004).

55 A. Pérez, I. Marchán, D. Svozil, J. Sponer, T. E. Cheatham, C. A. Laughton, and M. Orozco, “Refinement of the AMBER force field for nucleic acids: Improving the description of *α*/*γ* conformers,” Biophys. J. 92, 3817–3829 (2007).

56 P. Banáš, D. Hollas, M. Zgarbová, P. Jurečka, M. Orozco, T. E. Cheatham, J. Šponer, and M. Otyepka, “Performance of molecular mechanics force fields for RNA simulations: Stability of UUCG and GNRA hairpins,” J. Chem. Theory Comput. 6, 3836–3849 (2010).

57 M. Zgarbová, M. Otyepka, J. Šponer, A. Mládek, P. Banáš, T. E. Cheatham, and P. Jurečka, “Refinement of the Cornell et al. Nucleic acids force field based on reference quantum chemical calculations of glycosidic torsion profiles,” J. Chem. Theory Comput. 7, 2886–2902 (2011).

58 I.-C. Yeh and G. Hummer, “System-Size Dependence of Diffusion Coefficients and Viscosities from Molecular Dynamics Simulations with Periodic Boundary Conditions,” J. Phys. Chem. B 108, 15873–15879 (2004).

59 R. K. Sigel and H. Sigel, “A stability concept for metal ion coordination to single-stranded nucleic acids and affinities of individual sites,” Acc. Chem. Res. 43, 974–984 (2010).

60 Y. Marcus, “Ionic Radii in Aqueous Solutions,” Chem. Rev. 88, 1475–1498 (1988).

61 J. Neely and R. Connick, “Rate of Water Exchange from Hydrated Magnesium Ion,” J. Am. Chem. Soc. 92, 3476–3478 (1970).

62 A. Bleuzen, P.-A. Pittet, L. Helm, and A. E. Merbach, “Water exchange on magnesium(II) in aqueous solution: a variable temperature and pressure 17O NMR study,” Magn. Reson. Chem. 35, 765–773 (1997).

63 R. A. Robinson and R. H. Stokes, Electrolyte Solutions, 2nd ed. (Dover, New York, 2002).

64 F. Leonarski, L. D’Ascenzo, and P. Auffinger, “Mg^2+^ ions: Do they bind to nucleobase nitrogens?” Nucleic Acids Res. 45, 987–1004 (2017).

65 M. D. Tissandier, K. A. Cowen, W. Y. Feng, E. Gundlach, M. H. Cohen, A. D. Earhart, J. V. Coe, and T. R. Tuttle, “The proton’s absolute aqueous enthalpy and Gibbs free energy of solvation from cluster-ion solvation data,” J. Phys. Chem. A 102, 7787–7794 (1998).

66 T. L. Beck, “The influence of water interfacial potentials on ion hydration in bulk water and near interfaces,” Chem. Phys. Lett. 561-562, 1–13 (2013).

67 P. Loche, P. Steinbrunner, S. Friedowitz, R. R. Netz, and D. J. Bonthuis, “Transferable Ion Force Fields in Water from a Simultaneous Optimization of Ion Solvation and Ion–Ion Interaction,” J. Phys. Chem. B 125, 8581–8587 (2021).

68 S. Cruz-León, W. Vanderlinden, P. Müller, T. Forster, G. Staudt, Y. Lin, J. Lipfert, and N. Schwierz, “Twisting DNA by Salt,” bioRxiv2021.07.14.452306, 21–26 (2021).

