## Supplementary Information for "Magnesium Force Fields for OPC Water with Accurate Solvation, Ion-Binding, and Water-Exchange Properties: Successful Transfer from SPC/E"

#### Contents

**S1 Parameters of different water models** **S2**

**S2 Parameters of different $Mg^{2+}$ force fields** **S2**

**S3 Simulation setup** **S3**

**S4 Solvation free energy** **S4**

**S5 Exchange rates from long straight forward simulations** **S6**

**S6 Activity derivative of $MgCl_2$ solutions** **S9**

**S7 Binding affinity to DMP** **S10**

S7.1 Integration of potentials of mean force . . . . . . . . . . . . . . . . . . . . . . . . . . . . . . S11

S7.2 Alchemical transformation calculations . . . . . . . . . . . . . . . . . . . . . . . . . . . . . . S12

**S8 Bibliography** **S13**

### S1 Parameters of different water models

Table S1: Parameters of different water models. The 3-site waters (TIP3P, SPC/E, TIP3P-fb) carry their negative partial charge ($q_{\mathrm{Ow/Mw}}$) on the oxygen Ow. For the 4-site waters (TIP4P/2005, TIP4P-Ew, TIP4P-D, and OPC), it is located on the dummy atom Mw. $q_{\mathrm{H}}$ is the partial charge of the hydrogen atoms and $\sigma_{\mathrm{Ow}}$ and $\varepsilon_{\mathrm{Ow}}$ the Lennard-Jones parameters of the water oxygen. The length of the oxygen hydrogen bond is $l$, the angle between hydrogen, oxygen, and hydrogen is $\theta$. The distance between Ow and Mw is given for the 4-site waters as $z$.

| | $q_{\mathrm{Ow/Mw}}$ [e] | $q_{\mathrm{H}}$ [e] | $\sigma_{\mathrm{Ow}}$ [nm] | $\varepsilon_{\mathrm{Ow}}$ [kJ/mol] | $l$ [nm] | $\theta$ [°] | $z$ [nm] |
|---|---|---|---|---|---|---|---|
| TIP3P | -0.834 | 0.417 | 0.315061 | 0.636386 | 0.09572 | 104.52 | n.a. |
| SPC/E | -0.8476 | 0.4238 | 0.3166 | 0.650 | 0.1 | 109.47 | n.a. |
| TIP3P-fb | -0.848448 | 0.424224 | 0.31779(6) | 0.65214(4) | 0.10118 | 108.1484(4) | n.a. |
| TIP4P/2005 | -1.1128 | 0.5564 | 0.31589 | 0.774898 | 0.09572 | 104.52 | 0.01546 |
| TIP4P-Ew | -1.04844 | 0.52422 | 0.316435 | 0.680946 | 0.09572 | 104.52 | 0.0125 |
| TIP4P-D | -1.16 | 0.58 | 0.316508 | 0.936256 | 0.09572 | 104.52 | 0.0125 |
| **OPC** | **-1.3582** | **0.6791** | **0.316655** | **0.89036** | **0.08724** | **103.6** | **0.01594** |
| exp. (gas) | n.a. | n.a. | n.a. | n.a. | 0.09572 | 104.54 | n.a |

### S2 Parameters of different $Mg^{2+}$ force fields

Table S2: Parameters after transferring to OPC. *MicroMg* and *nanoMg* parameters, optimized in TIP3P, SPC/E, TIP3P-fb, TIP4P/2005, TIP4P-Ew, or TIP4P-D, and adapted to OPC. The OPC-optimized Li-Merz parameters are also listed. $\sigma_{\mathrm{ii}}$, $\varepsilon_{\mathrm{ii}}$, $\sigma_{\mathrm{io}}$, $\varepsilon_{\mathrm{io}}$ are the ion-ion and ion-water LJ parameters adapted to OPC (see eq 2 in the main text). $\lambda_\sigma^{\mathrm{X}}$ and $\lambda_\varepsilon^{\mathrm{X}}$ are the scaling factors for the Lorentz-Berthelot combination rules (eq 3 in the main text) for the interaction with $Cl^-$ or the RNA atoms. The 12-6 and 12-6-4 parameters by Li et al. are taken from ref.[1] The additional parameter of the 12-6-4 potential for the $1/r_{\mathrm{ij}}^4$ term is $C_4 = 0.0531368$ kJ nm$^4$/mol.

| | $\sigma_{\mathrm{ii}}$ [nm] | $\varepsilon_{\mathrm{ii}}$ [kJ/mol] | $\sigma_{\mathrm{io}}$ [nm] | $\varepsilon_{\mathrm{io}}$ [kJ/mol] | $\lambda_\sigma^{\mathrm{Cl}}$ | $\lambda_\varepsilon^{\mathrm{Cl}}$ | $\sigma_{\mathrm{MgCl}}$ [nm] | $\varepsilon_{\mathrm{MgCl}}$ [kJ/mol] | $\lambda_\sigma^{\mathrm{RNA}}$ | $\lambda_\varepsilon^{\mathrm{RNA}}$ | $\sigma_{\mathrm{MgOP}}$ [nm] | $\varepsilon_{\mathrm{MgOP}}$ [kJ/mol] |
|---|---|---|---|---|---|---|---|---|---|---|---|---|
| TIP3P | | | | | | | | | | | | |
| *microMg* | 0.1019 | 235.80 | 0.2093 | 14.49 | 1.80 | 0.1 | 0.4878 | 0.8181 | 1.1375 | 0.3200 | 0.2262 | 4.6061 |
| *nanoMg* | 0.1025 | 389.80 | 0.2096 | 18.63 | 1.80 | 0.1 | 0.4884 | 1.0518 | 1.1435 | 0.2500 | 0.2277 | 4.6266 |
| SPC/E | | | | | | | | | | | | |
| *microMg* | 0.1036 | 290.58 | 0.2101 | 16.08 | 1.59 | 0.1 | 0.4318 | 1.0989 | 1.1019 | 0.4856 | 0.2202 | 7.7589 |
| *nanoMg* | 0.1046 | 470.70 | 0.2106 | 20.47 | 1.59 | 0.1 | 0.4325 | 1.3986 | 1.1107 | 0.3300 | 0.2225 | 6.7111 |
| TIP3P-fb | | | | | | | | | | | | |
| *microMg* | 0.1032 | 311.38 | 0.2099 | 16.65 | 1.59 | 0.1 | 0.4743 | 0.3983 | 1.0957 | 0.4913 | 0.2187 | 8.1266 |
| *nanoMg* | 0.1034 | 380.38 | 0.2100 | 18.40 | 1.59 | 0.1 | 0.4744 | 0.4403 | 1.1002 | 0.4172 | 0.2197 | 7.6266 |
| TIP4P/2005 | | | | | | | | | | | | |
| *microMg* | 0.0901 | 712.67 | 0.2034 | 25.19 | 1.59 | 0.1 | 0.4719 | 0.5529 | 1.1484 | 0.2648 | 0.2217 | 6.6266 |
| *nanoMg* | 0.0913 | 774.62 | 0.2040 | 26.26 | 1.59 | 0.1 | 0.4728 | 0.5764 | 1.1345 | 0.2923 | 0.2217 | 6.6266 |
| TIP4P-D | | | | | | | | | | | | |
| *microMg* | 0.0960 | 621.50 | 0.2063 | 23.52 | 1.59 | 0.1 | 0.4684 | 0.5622 | 1.1456 | 0.2778 | 0.2217 | 6.6266 |
| *nanoMg* | 0.0970 | 680.01 | 0.2068 | 24.61 | 1.59 | 0.1 | 0.4697 | 0.6096 | 1.1489 | 0.2371 | 0.2232 | 6.1266 |
| TIP4P-Ew | | | | | | | | | | | | |
| *microMg* | 0.0910 | 647.63 | 0.20380 | 24.01 | 1.59 | 0.1 | 0.4811 | 0.4697 | 1.3255 | 0.3264 | 0.2197 | 7.6266 |
| *nanoMg* | 0.0926 | 760.06 | 0.2046 | 26.01 | 1.59 | 0.1 | 0.4819 | 0.4913 | 1.1231 | 0.2834 | 0.2207 | 6.9266 |
| OPC | | | | | | | | | | | | |
| Li-Merz (12-6) | 0.2208 | 0.00864 | 0.2687 | 0.08769 | n.a. | n.a. | 0.3019 | 0.1373 | n.a. | n.a. | 0.2584 | 0.0871 |
| Li-Merz (12-6-4) | 0.2503 | 0.06916 | 0.2835 | 0.24815 | n.a. | n.a. | 0.3161 | 0.3863 | n.a. | n.a. | 0.2732 | 0.2465 |

### S3 Simulation setup

In all simulations, the short-range Coulomb and Lennard-Jones interactions were cut off at 1.2 nm. For long-range electrostatic forces, we employed tin-foil periodic boundary conditions, the particle-mesh Ewald method,[2] and a Fourier spacing of 0.12 nm with grid interpolation up to order 4. With the rigid OPC water model (Table S1), we used a 2 fs timestep. Employing AMBER, we used SETTLE.[3] For simulations with GROMACS, we used LINCS[4] to constrain hydrogen bonds in simulations containing DMP. Frames were written every 2 ps if not mentioned otherwise. Energy minimization was performed by steepest decent, followed by two equilibration runs of 0.5 ns in NVT and 1 ns in NPT. These simulations were performed at 300 K and 1 bar, using thermostat and barostat of Berendsen.[5] During the equilibration, we used position restraints on all ions to make sure that the hydration shells form properly and prevent the artificial formation of inner-sphere $Mg^{2+}$-$Cl^-$ ion pairs (see radial distribution function in Figure S2). In the production runs, these restraints were released. For the calculation of solvation free energy $\Delta G_{\mathrm{solv}}$, radius of the first hydration shell $R_1$, and coordination number $n_1$, we employed the thermostat and barostat of Berendsen[5] with $\tau = 0.1$ and $\tau_{\mathrm{p}} = 1.0$ to ensure a temperature of 300 K and an atmospheric pressure close to 1 bar. In all other simulation, the velocity rescaling thermostat by Bussi et al.[6] with $\tau = 0.1$ was used. For the NPT simulations (Table S3), the Parrinello-Rahman barostat[7] with $\tau_{\mathrm{p}} = 5.0$ was applied. We used the same setups (Table S3) for simulations with GROMACS and AMBER and employed Lorentz-Berthelot combination rules (eq 2, main text) in all simulations.

Table S3: Simulation setups. In 'Phys. property', the physical properties are listed that we obtained from the respective simulation using the respective 'Method' (FEP: free energy perturbation, unbiased: straight forward simulations without additional biases, US: umbrella sampling). 'System' lists all particles of the respective simulation and 'Duration' their duration (products show the number of window times their duration). '$L$' indicates the simulation box size for (cu) cubic and (do) rhombic dodecahedron box shapes. 'Ensemble' expresses if an canonical ensemble (with constant number of particles $N$, constant volume $V$, and constant temperature $T$), or an isobaric-isothermal ensemble (with constant pressure $P$) was used. GMX and AMBER specify the setup used for the 12-6 based (GMX) or the 12-6-4 based parameters (AMBER).

| Phys. property | Method | System | Duration | $L$ | Ensemble |
|---|---|---|---|---|---|
| $\Delta G_{\mathrm{solv}}$, $R_1$, $n_1$ | FEP | 1 $Mg^{2+}$, 506 water | 40 × 1 ns | 2.5 nm (cu) | NPT |
| $k$ | unbiased (AMBER) | 39 $Mg^{2+}$, 78 $Cl^-$, 2048 water (1 M) | 4 × 500 ns | 4 nm (cu) | NPT |
| $a_{\mathrm{cc}}$, $k$ | unbiased (GMX) | 39 $Mg^{2+}$, 78 $Cl^-$, 2055 water (1 M) | 2 μs | 4 nm (cu) | NPT |
| $a_{\mathrm{cc}}$ | unbiased | 73 $Mg^{2+}$, 146 $Cl^-$, 1909 water (2 M) | 150 ns | 4 nm (cu) | NPT |
| $a_{\mathrm{cc}}$ | unbiased | 20 $Mg^{2+}$, 40 $Cl^-$, 2135 water (0.5 M) | 150 ns | 4 nm (cu) | NPT |
| $a_{\mathrm{cc}}$ | unbiased | 10 $Mg^{2+}$, 20 $Cl^-$, 2171 water (0.25 M) | 150 ns | 4 nm (cu) | NPT |
| $F(r_{\mathrm{MgOw}})$ | US (GMX) | 1 $Mg^{2+}$, 505 water | 68 × 3 ns | 2.5 nm (cu) | NPT |
| $F(r_{\mathrm{MgOw}})$ | US (AMBER) | 1 $Mg^{2+}$, 2 $Cl^-$, 506 water | 68 × 3 ns | 2.5 nm (cu) | NPT |
| $\Delta G^0_{\mathrm{b}}$ | FEP | 1 DMP, 1 $Mg^{2+}$, 1 $Cl^-$, 1536 water | 20 × 9 ns | 4 nm (do) | NPT |
| $\Delta G^0_{\mathrm{b}}$, $F(r_{\mathrm{MgOP}})$ | US | 1 DMP, 1 $Mg^{2+}$, 1 $Cl^-$, 1536 water | 67 × 20 ns | 4 nm (do) | NVT |
| $\Delta G^0_{\mathrm{ref}}$, $F(r_{\mathrm{ref-OP}})$ | US | 1 DMP, 1 ref atom, 1 $Cl^-$, 1536 water | 67 × 20 ns | 4 nm (do) | NVT |

#### S4 Solvation free energy

Solvation free energies were obtained for the transferred parameters of the neutral $MgCl_2$ ion pair with free energy perturbation (FEP) and Bennett's acceptance ratio (BAR) method.[8] The statistical uncertainty is about 1 kJ/mol. The perturbation calculation was performed for each parameter set over 40 evenly spaced intermediate states, simulated for 1 ns, first (states $0-19$) creating neutral Lennard-Jones particles, and second (states $20-39$) increasing the positive net charge of the particle until it reaches 2. To avoid divergences, we employed soft-core potentials. In the steps $0-19$, $A$ corresponds to the fully uncoupled system and $B$ characterizes a system containing the neutral Lennard-Jones particle. In the steps $20-39$, $A$ corresponds to the neutral LJ particle and $B$ to the fully coupled system.

We excluded the first 200 ps from each intermediate state for equilibration. We applied several correction factors to be able to directly compare experiment and simulations. To account for finite size effects, we employed[9]

$$\Delta G_{\rm fs} = \frac{z^2 N_A e^2}{4\pi\epsilon_0}\left[-\frac{\zeta_{\rm ew}}{2\epsilon_r} + \left(1-\frac{1}{\epsilon_r}\right)\left(\frac{2\pi R_1^2}{3L^3} - \frac{8\pi^2 R_1^5}{45L^6}\right)\right] , \tag{S1}$$

where $z$ is the valency, $N_A$ Avogardo's number, $e$ the unit charge, $\epsilon_0$ the vacuum permittivity, $R_1$ the first peak of the ion-water radial distribution function, $\zeta_{\rm ew} = -2.837297/L$ is the Wigner potential with $L$ being the edge length of the simulation box in nm.[9,10] $\epsilon_r$ is the relative dielectric constant of the different water models. We used $\epsilon_r = 78.4$ for OPC water.[11]

To account for the compression of ideal gas with $p_0 = 1$ bar to the pressure of an ideal solution at a density of 1 mol/l with $p_1 = 26.4$ bar, we used the correction term[12]

$$\Delta G_{\rm press} = N_{\rm A} k_{\rm B} T \ln(p_1/p_0) = 7.9 \text{ kJ/mol} . \tag{S2}$$

Finally, in the typical experimental setup, ions have to pass the air-water interface to enter the aqueous phase, which is accounted for in simulations with the correction term[12]

$$\Delta G_{\rm surf} = N_A z \times e\phi_{\rm surf} = -z \times 50.8 \text{ kJ/mol} , \tag{S3}$$

where we chose the surface potential to be $\phi_{\rm surf} = -0.527$ V.[13,14] This value closely matches the experimental estimation of -0.50 V.[15] Including all correction terms, the solvation free energy of a single ion $X$ is obtained from,

$$\Delta G^X_{\rm solv} = \Delta G^X_{\rm sim} + \Delta G_{\rm fs} + \Delta G_{\rm surf} + \Delta G_{\rm press} . \tag{S4}$$

For the neutral ion pair it is given by

$$\Delta G_{\text{solv}} = \Delta G_{\text{solv}}^{\text{Mg}^{2+}} + z \times \Delta G_{\text{solv}}^{\text{Cl}^-} \ . \tag{S5}$$

Note that for neutral ion pairs the surface term cancels in the calulation of the solvation free energy $\Delta G_{\text{solv}}$. However, for solvation free energy of a single $Cl^-$ ion, $\Delta G_{\text{solv}}^{\text{Cl}^-}$, it needs to be considered in order to compare to the experimental value which includes the interfacial crossing.[15,16] The calculated values are listed in Tables S4 and S5.

Table S4: Single-ion properties after transferring the *microMg* and *nanoMg* parameters to OPC. $\Delta G_{\text{solv}}$, $R_1$, and $n_1$ are the solvation free energy of the neutral $MgCl_2$ ion-pair, the radius of the first hydration shell, and the coordination number of the first hydration shell in OPC. Properties for the Li-Merz parameters were taken from ref.,[1] where the solvation free energy was reported for a single $Mg^{2+}$ ion. To compare with $\Delta G_{\text{solv}}$ for neutral ion pairs, we added the experimental $Cl^-$ value from Marcus[17] $\Delta G_{\text{solv}}^{\text{Cl}^-}$= 347 kJ/mol.

| | $\Delta G_{\text{solv}}$ [kJ/mol] | $R_1$ [nm] | $n_1$ |
|---|---|---|---|
| *microMg*(TIP3P) | -2546.6 ± 1 | 0.211 ± 0.004 | 6 |
| *nanoMg*(TIP3P) | -2553.6 ± 1 | 0.214 ± 0.004 | 6 |
| *microMg*(SPC/E) | -2628.7 ± 1 | 0.212 ± 0.004 | 6 |
| *nanoMg*(SPC/E) | -2535.9 ± 1 | 0.215 ± 0.004 | 6 |
| *microMg*(TIP3P-fb) | -2521.3 ± 1 | 0.212 ± 0.004 | 6 |
| *nanoMg*(TIP3P-fb) | -2525.5 ± 1 | 0.214 ± 0.004 | 6 |
| *microMg*(TIP4P/2005) | -2597.8 ± 1 | 0.210 ± 0.004 | 6 |
| *nanoMg*(TIP4P/2005) | -2596.6 ± 1 | 0.211 ± 0.004 | 6 |
| *microMg*(TIP4P-D) | -2592.8 ± 1 | 0.210 ± 0.004 | 6 |
| *nanoMg*(TIP4P-D) | -2592.6 ± 1 | 0.211 ± 0.004 | 6 |
| *microMg*(TIP4P-Ew) | -2564.8 ± 1 | 0.212 ± 0.004 | 6 |
| *nanoMg*(TIP4P-Ew) | -2563.4 ± 1 | 0.214 ± 0.004 | 6 |
| 12-6 Li-Merz (OPC) | -2522.0 | 0.195 | 6 |
| 12-6-4 Li-Merz (OPC) | -2524.9 | 0.209 | 6 |
| exp. | -2532[17] | 0.209 ± 0.004[18] | 6[18] |

Table S5: Parameters and single-ion properties for the Cl- parameters transferred to OPC. $\sigma_{ii}$, $\varepsilon_{ii}$, $\sigma_{io}$, $\varepsilon_{io}$ are the ion-ion and ion-water LJ parameters adapted to OPC (see eq 2 in the main text). $\Delta G_{solv}^{Cl^-}$ and $R_1$ are the solvation free energy and the radius of its first hydration shell in OPC. The parameters and properties for OPC-optimized 12-6 and 12-6-4 Sengupta-Merz parameters were taken from ref.[19] The additional parameter of the 12-6-4 potential for the $1/r_{ij}^4$ term is $C_4$ = -0.0288696 kJ nm$^4$/mol.

| | $\sigma_{ii}$ [nm] | $\varepsilon_{ii}$ [kJ/mol] | $\sigma_{io}$ [nm] | $\varepsilon_{io}$ [kJ/mol] | $\Delta G_{solv}^{Cl^-}$ [kJ/mol] | $R_1^{Cl^-}$ [nm] |
|---|---|---|---|---|---|---|
| Mamatkulov-Schwierz(TIP3P) | 0.440939 | 0.283829 | 0.378797 | 0.502703 | -317.9 ± 1 | 0.319 |
| Smith-Dang(SPC/E) | 0.439443 | 0.415598 | 0.378049 | 0.608302 | -311.3 ± 1 | 0.326 |
| Grotz-Schwierz(TIP3P-fb) | 0.493358 | 0.050960 | 0.405007 | 0.213009 | -306.1 ± 1 | 0.320 |
| Grotz-Schwierz(TIP4P/2005) | 0.503464 | 0.042887 | 0.410060 | 0.195409 | -300.2 ± 1 | 0.323 |
| Grotz-Schwierz(TIP4P-Ew) | 0.498225 | 0.048805 | 0.407440 | 0.208456 | -301.3 ± 1 | 0.323 |
| Grotz-Schwierz(TIP4P-D) | 0.509112 | 0.035496 | 0.412884 | 0.177776 | -300.0 ± 1 | 0.323 |
| 12-6 Sengupta-Merz (OPC) | 0.420504 | 0.069159 | 0.368580 | 0.087690 | -373.2 | 0.342 |
| 12-6-4 Sengupta-Merz (OPC) | 0.381839 | 2.15745 | 0.349247 | 1.38597 | -374.5 | 0.328 |
| exp. | | | | | -304.2[17] | 0.319 ± 0.007[18] |

### S5 Exchange rates from long straight forward simulations

The most popular theory to calculate reaction rates is transition state theory (TST).[20,21] TST gives an accurate estimate of the rate for simple systems, in which the reaction coordinate is exactly known. However, in many body systems, complex processes such as the exchange of waters in the first hydration shell can cause TST to fail due to the violation of the non-recrossing hypothesis,[22] one of the fundamentals of the theory. We therefore used long straight forward simulations to obtain the rate constant. The rate constant $k$ of the exchange of water from the first hydration shell of $Mg^{2+}$ is defined by[23]

$$\text{rate} = 6 \cdot k \cdot [\text{Mg}(\text{H}_2\text{O})_6^{2+}] \ , \tag{S6}$$

where 6 is the coordination number of the first hydration shell and $[\text{Mg}(\text{H}_2\text{O})_6^{2+}]$ is the concentration of hexa-coordinated $Mg^{2+}$ ions.

In this work, the water exchange rate constant $k$ was calculated by counting the total number of transitions that were observed within 2 $\mu s$ long trajectories of a 1 M $MgCl_2$ solution. We count the transitions $N$ of exchanges between the first and second hydration shell (the exchange from first to second hydration shell and the reverse transition are counted as individual events). The water exchange rate constant $k$ is given by

$$k = \frac{1}{N_{\text{H}_2\text{O}}} \cdot \frac{N}{2 \cdot t_\text{B}} \ , \tag{S7}$$

where $N_{\text{H}_2\text{O}}$ is the number of water molecules within the simulation box. $t_\text{B} = N_\text{Mg} \cdot p_\text{B} \cdot t_\text{sim}$ is the cumulative time the water molecule spends in the first hydration shell of any $Mg^{2+}$ ion. $N_\text{Mg}$ is the number of $Mg^{2+}$

ions in the simulation box, $p_B = 6/(N_{H_2O} - 6)$ is the probability of water to be in the first hydration shell and $t_{sim}$ is the total simulation time. Errors were calculated from block averaging by dividing the trajectory into 2 blocks. The rate constants obtained in this work are shown in Table S6.

Table S6: Properties of water exchange from simulations and experiments. Number of transitions $N$ in 2 $\mu$s for different $Mg^{2+}$ parameters in 1 M $MgCl_2$ solutions. The experimental value for $N$ is obtained from eq S7 using the experimental value for the rate from refs.[23,24] The errors for $N$ and $k$ are obtained from block averaging. $\Delta F^\dagger$ is the free energy difference between the first minimum and the barrier top in the free energy profiles (Figure S1).

| | $N$ | $k$ [$s^{-1}$] | $\Delta F^\dagger$ [$k_B$T] |
|---|---|---|---|
| *microMg*(TIP3P) | 8 ± 4 | (1.28 ± 0.9) · $10^4$ | 19.8 |
| *nanoMg*(TIP3P) | 806 ± 26 | (1.29 ± 0.1) · $10^6$ | 15.9 |
| *microMg*(SPC/E) | 100 ± 20 | (1.22 ± 0.5) · $10^5$ | 17.3 |
| *nanoMg*(SPC/E) | 17,869 ± 341 | (2.86 ± 0.07) · $10^7$ | 12.7 |
| *microMg*(TIP3P-fb) | 142 ± 6 | (1.84 ± 0.08) · $10^5$ | 17.4 |
| *nanoMg*(TIP3P-fb) | 935 ± 13 | (1.50 ± 0.1) · $10^6$ | 15.3 |
| 12-6 Li-Merz (OPC) | 12 ± 12 | (1.92 ± 1.92) · $10^4$ | 15.1 |
| 12-6-4 Li-Merz (OPC) | 0 ± 0 | n.a. | 20.5 |
| exp. | 496[24] , 628[23] | 5.3 · $10^5$ from[23] , 6.7 · $10^5$ from[24] | n.a. |

One-dimensional free energy profiles $F(R)$ were obtained from umbrella sampling[25,26] as a function of the distance between $Mg^{2+}$ and the oxygen atom of the leaving water molecule for the 12-6 type force fields with GROMACS[27] (version 2020). The profile with the 12-6-4 type force field in OPC water was computed using AMBER[28] (version 2018) and PLUMED[29] (version 2.5). Force constants and window spacing were $k = 400,000$ kJ/(mol nm$^2$) and 0.005 nm [$0.17 \leq R_{MgOx} < 0.4$ nm] and $k = 100,000$ kJ/(mol nm$^2$) and 0.01 nm [$0.4 \leq R_{MgOx} < 0.6$ nm]. Frames for the analysis were considered every 0.5 ps. We used the weighted histogram analysis method (WHAM)[30] to combine the individual umbrella windows with a bin width of $5.4 \cdot 10^{-4}$ nm.

The free energy profiles along the distance between $Mg^{2+}$ and one of the two non-bridging phosphate oxygens of the dimethylphosphate (DMP) (Figure 3C, main text) were obtained with force constants and window spacing of $k = 300,000$ kJ/(mol nm$^2$) and 0.0075 nm [$0.15 \leq R_{MgOP} < 0.525$ nm] and $k = 5,000$ kJ/(mol nm$^2$) and 0.02 nm [$0.525 \leq R_{MgOP} < 0.885$ nm], respectively, using GROMACS[27] (version 2020) and PLUMED.[29] An additional bias was applied to avoid artificial contacts with DMP atoms other than the selected phosphate oxygen (see ref.[31] for further details). Frames for the analysis were considered every 0.5 ps. For WHAM a bin width of $9.2 \cdot 10^{-4}$ nm was considered.

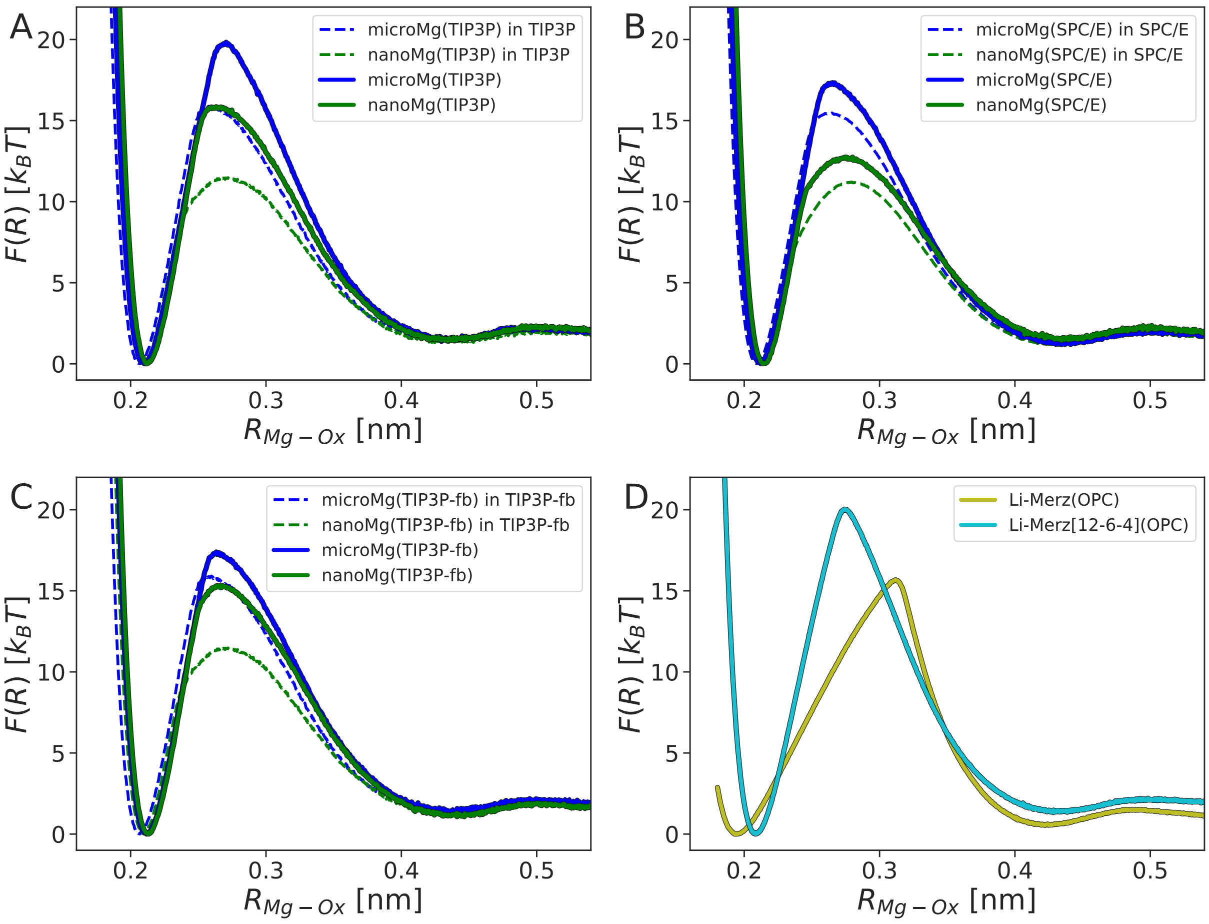


Figure S1: One-dimension free energy profiles along the distance between $Mg^{2+}$ and a water oxygen. (A) TIP3P-optimized $Mg^{2+}$ parameters in OPC. (B) SPC/E-optimized $Mg^{2+}$ parameters in OPC. (C) TIP3P-fb-optimized $Mg^{2+}$ parameters in OPC. (D) OPC-optimized 12-6 and 12-6-4 Li-Merz parameters. In (A-C), the results in the original water model are shown as dashed lines. Note that the free energy barrier in the original water model is always lower compared to the barrier in OPC. In (D) the barrier for the 12-6 Li-Merz parameters is lower compared to the 12-6-4 parameters which explains the absence of exchange events (Table S6). Note that this observation is opposite to the result in TIP3P and SPC/E where the barrier for the 12-6-4 interaction potentials is lower compared to 12-6 potentials.[31,32]

#### S6 Activity derivative of $MgCl_2$ solutions

We used Kirkwood-Buff (KB) theory to obtain the activity derivative. The KB integrals have the following form[14,33,34]

$$G_{\mathrm{ij}} = 4\pi \int_0^\infty \left[g_{\mathrm{ij}}^{\mu \mathrm{VT}}(r_{\mathrm{ij}}) - 1\right] r_{\mathrm{ij}}^2 dr_{\mathrm{ij}} \ , \tag{S8}$$

where $g_{\mathrm{ij}}^{\mu \mathrm{VT}}(r_{\mathrm{ij}})$ is the radial distribution function of two species in the grand canonical ensemble, with $r_{\mathrm{ij}}$ being the center of mass distance between the two. Note that the simulation were done in the NPT ensemble. Therefore, we introduced a correction factor such that the radial distribution function used in the calculations of the KB integrals shows the correct asymptotic behavior at large distances. For divalent cations, the equations have the following for,[34,35]

$$G_{\mathrm{cc}} = \frac{1}{9}\left[G_{++} + 4(G_{--} + G_{+-})\right] \tag{S9}$$

and

$$G_{\mathrm{co}} = G_{\mathrm{oc}} = \frac{1}{3}G_{+\mathrm{o}} + \frac{2}{3}G_{-\mathrm{o}} \ , \tag{S10}$$

with $+$, $-$, and o denoting the cation, anion, and water oxygen, respectively. The derivative $a_{\mathrm{cc}}$ of the activity $a_{\mathrm{c}} = \rho_{\mathrm{c}} y_{\mathrm{c}}$ with activity coefficient $y_{\mathrm{c}}$ is defined via

$$a_{\mathrm{cc}} = \left(\frac{\partial \ln a_{\mathrm{c}}}{\partial \ln \rho_{\mathrm{c}}}\right)_{\mathrm{P,T}} = 1 - \left(\frac{\partial \ln y_{\mathrm{c}}}{\partial \ln \rho_{\mathrm{c}}}\right)_{\mathrm{P,T}} = \frac{1}{1 + \rho_{\mathrm{c}}(G_{\mathrm{cc}} - G_{\mathrm{co}})} \ , \tag{S11}$$

with respect to the natural logarithm of the number density $\rho_{\mathrm{c}}$.

Figure S2 shows the $Mg^{2+}$-$Cl^{-}$ radial distribution function $g_{\mathrm{MgCl}}(r_{\mathrm{MgCl}})$ for different force fields. From the position of the first peak of the radial distribution functions it becomes evident that no direct inner-sphere interactions between $Mg^{2+}$ and $Cl^{-}$ ions occur and that their interaction is always mediated by their hydration shells, as expected.

The activity derivatives obtained in this work are shown in Figure 3A in the main text.

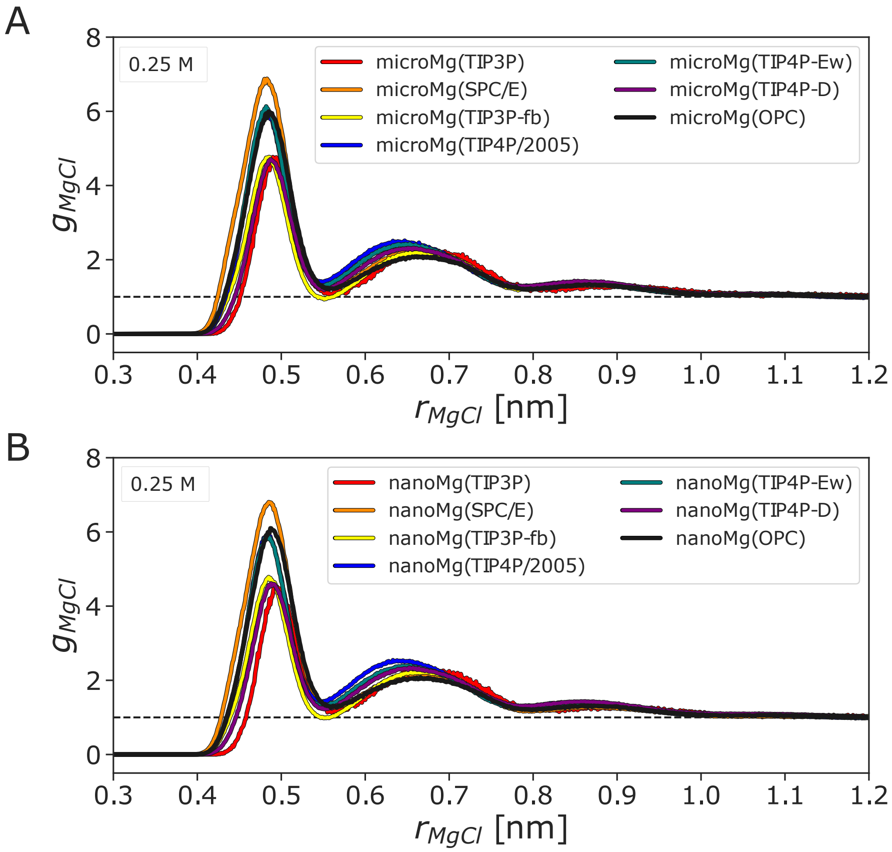


Figure S2: Radial distribution functions $g_{\mathrm{MgCl}}$ along the distance $r_{\mathrm{MgCl}}$ between $Mg^{2+}$ and $Cl^-$ with (A) *microMg* and (B) *nanoMg* for different water models. The dashed line indicates unity as asymptotically reached at large distances. Simulations were done at 0.25 M salt concentration.

#### S7 Binding affinity to DMP

The binding affinity $\Delta G_{\mathrm{b}}^{0}$ of the $Mg^{2+}$ ion toward one of the non-bridging phosphate oxygens of the backbone was considered. The RNA is mimicked by a dimethylphosphate (DMP). We calculated $\Delta G_{\mathrm{b}}^{0}$ by integrating the potential of mean force along the distance between $Mg^{2+}$ and one of the non-bridging phosphate oxygens of the DMP. For the final parameter set, a second method (alchemical transformation) was employed to validate the results.

#### S7.1 Integration of potentials of mean force

The binding affinity is obtained by integrating the potential of mean force $V^{\mathrm{PMF}}$ along the distance $r$ between the binding site and the ion,

$$\Delta G_{\mathrm{b}}^{0} = -k_{\mathrm{B}}T \cdot \ln\left(\frac{c^0}{[M]} \cdot \frac{\int_0^{\mathrm{r}^\dagger} r^2 e^{-\mathrm{V}^{\mathrm{PMF}}(\mathrm{r})/\mathrm{k_B T}} \mathrm{d}r}{\int_{\mathrm{r}^\dagger}^{\mathrm{r_L}} r^2 e^{-\mathrm{V}^{\mathrm{PMF}}(\mathrm{r})/\mathrm{k_B T}} \mathrm{d}r}\right), \tag{S12}$$

where $[M]$ is the ion concentration of the simulation box, $r_{\mathrm{L}}$ is the radius of a sphere that contains the same number of water molecules as the simulation box and $r^\dagger$ is the position of the maximum of the PMF. Note that both $[M]$ and $r_{\mathrm{L}}$ are dependent on the number of waters in the simulation such that eq S12 becomes independent of the box size used.

The potentials of mean force $V^{\mathrm{PMF}}(r)$ were obtained from free energy profiles $F(r)$ by applying a Jacobian correction

$$V^{\mathrm{PMF}}(r) = F(r) + 2k_{\mathrm{B}}T \ln r. \tag{S13}$$

Free energy profiles $F(R)$ were calculated with umbrella sampling[25,26] and the weighted histogram analysis method (WHAM)[30] (see Section S5) and are shown in Figure 3C of the main text and in Figure S3). The binding affinities $\Delta G_{\mathrm{b}}^{0}$ obtained from integration of $V^{\mathrm{PMF}}(r)$ are listed in Table S7.

Table S7: Binding affinity $\Delta G_{\mathrm{b}}^{0}$ toward one of the non-bridging phosphate oxygens of a dimethylphosphate. $R_{\mathrm{b}}$ is the distance in the bound state and $\Delta F^\dagger$ the free energy difference between the first minimum in the free energy profile and the barrier top (Figure 3C of main text and Figure S3).

| | $\Delta G_{\mathrm{b}}^{0}$ [$\mathrm{k_B T}$] | $R_{\mathrm{b}}$ [nm] | $\Delta F^\dagger$ [$\mathrm{k_B T}$] |
|---|---|---|---|
| *microMg*(TIP3P) | 0.659 ± 0.6 | 0.206 | 18.7 |
| *nanoMg*(TIP3P) | 0.969 ± 0.5 | 0.207 | 15.0 |
| *microMg*(SPC/E) | -0.818 ± 0.5 | 0.207 | 15.1 |
| *nanoMg*(SPC/E) | -0.969 ± 0.3 | 0.207 | 12.1 |
| *microMg*(TIP3P-fb) | -2.209 ± 0.2 | 0.206 | 15.0 |
| *nanoMg*(TIP3P-fb) | -1.724 ± 0.3 | 0.206 | 13.7 |
| exp. | -1.036[36] | 0.206 - 0.208[37] | n.a. |

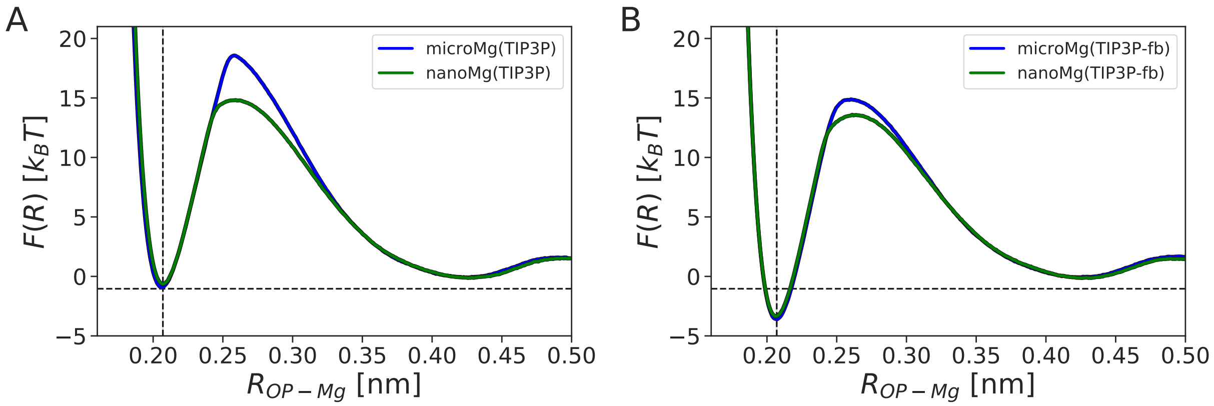


Figure S3: One-dimension free energy profile along the distance between $Mg^{2+}$ and one of the non-bridging phosphate oxygens of the DMP for $Mg^{2+}$ obtained in OPC water with parameters transferred from (A) TIP3P and (B) TIP3P-fb water. Dashed horizontal and vertical lines correspond to the experimental binding affinity[36] and the experimental binding distance[37] between the ion and its binding site in the bound state.

#### S7.2 Alchemical transformation calculations

As validation, we obtained the binding affinities of the SPC/E-optimized $Mg^{2+}$ parameters from free energy perturbation using alchemical transformation calculations[38,39] and MBAR.[40] A reference ion of the same valency was used. The ion of interest is alchemically transformed into the reference ion by gradually switching its van der Waals interactions, both in bulk ($\Delta G^{\mathrm{solv}}_{\mathrm{ion}\to\mathrm{ref}}$) as well as at the binding site ($\Delta G^{\mathrm{BS}}_{\mathrm{ref}\to\mathrm{ion}}$)

$$\Delta G^0_{\mathrm{b}} = \Delta G^{\mathrm{solv}}_{\mathrm{ion}\to\mathrm{ref}} + \Delta G^0_{\mathrm{ref}} + \Delta G^{\mathrm{BS}}_{\mathrm{ref}\to\mathrm{ion}} \tag{S14}$$

where $\Delta G^0_{\mathrm{ref}}$ is the binding affinity of the reference ion.

Convergence can be checked by transforming in opposite directions (*i.e.*, forward and backward, see Tables S8, S9). For the reference ion, we used the parameters for $Ca^{2+}$ obtained in TIP3P.[41] The binding affinity of $Ca^{2+}$ in OPC for eq S14 was $\Delta G^0_{\mathrm{ref}}$ = -5.153 $\mathrm{k_BT}$ (obtained from integrating the potential of mean force).

For SPC/E-optimized $Mg^{2+}$ in OPC, the binding affinities from the different methods are $0.8 \pm 0.5$ $\mathrm{k_BT}$, $0.5 \pm 0.4$ $\mathrm{k_BT}$ and $0.5 \pm 0.4$ $\mathrm{k_BT}$ and agree within error. Additional values are provided in Table S7, Table S8, and Table S9.

Table S8: Binding affinity $\Delta G_{\mathrm{b}}^{0}$ obtained from forward alchemical transformation. The values given for $\Delta G_{\mathrm{Mg^{2+}\to ref}}^{\mathrm{solv}}$ and $\Delta G_{\mathrm{ref\to Mg^{2+}}}^{\mathrm{bind}}$ are obtained from block averaging for 3 blocks of 3 ns long windows each. The binding affinity for the reference ion was $\Delta G_{\mathrm{ref}}^{0}$ = -5.153 $k_BT$.

| | $\Delta G_{\mathrm{b}}^{0}$ [$k_B$T] | $\Delta G_{\mathrm{Mg^{2+}\to ref}}^{\mathrm{solv}}$ [$k_B$T] | $\Delta G_{\mathrm{ref\to Mg^{2+}}}^{\mathrm{bind}}$ [$k_B$T] |
|---|---|---|---|
| *microMg*(SPC/E) | -0.535 ± 0.4 | 137.874 ± 0.003 | -133.256 ± 0.003 |
| *nanoMg*(SPC/E) | -0.205 ± 0.4 | 140.714 ± 0.003 | -135.765 ± 0.003 |
| exp. | -1.036[36] | n.a. | n.a. |

Table S9: Binding affinity $\Delta G_{\mathrm{b}}^{0}$ obtained from backward alchemical transformation. The values given for $\Delta G_{\mathrm{ref\to Mg^{2+}}}^{\mathrm{solv}}$ and $\Delta G_{\mathrm{Mg^{2+}\to ref}}^{\mathrm{bind}}$ are obtained from block averaging for 3 blocks of 3 ns long windows each. The binding affinity for the reference ion was $\Delta G_{\mathrm{ref}}^{0}$ = -5.153 $k_B$T.

| | $\Delta G_{\mathrm{b}}^{0}$ [$k_B$T] | $\Delta G_{\mathrm{ref\to Mg^{2+}}}^{\mathrm{solv}}$ [$k_B$T] | $\Delta G_{\mathrm{Mg^{2+}\to ref}}^{\mathrm{bind}}$ [$k_B$T] |
|---|---|---|---|
| *microMg*(SPC/E) | -0.458 ± 0.4 | -137.942 ± 0.003 | 133.246 ± 0.003 |
| *nanoMg*(SPC/E) | -0.333 ± 0.4 | -140.675 ± 0.003 | 135.855 ± 0.003 |
| exp. | -1.036[36] | n.a. | n.a. |

#### S8 Bibliography


(1) Li, Z.; Song, L. F.; Li, P.; Merz, K. M. Systematic Parametrization of Divalent Metal Ions for the OPC3, OPC, TIP3P-FB, and TIP4P-FB Water Models. *J. Chem. Theory Comput.* **2020**, *16*, 4429–4442.

(2) Darden, T.; York, D.; Pedersen, L. Particle mesh Ewald: An N x log(N) method for Ewald sums in large systems. *J. Chem. Phys.* **1993**, *98*, 10089–10092.

(3) Ryckaert, J. P.; Ciccotti, G.; Berendsen, H. J. Numerical integration of the cartesian equations of motion of a system with constraints: molecular dynamics of n-alkanes. *J. Comp. Phys.* **1977**, *23*, 327–341.

(4) Hess, B.; Bekker, H.; Berendsen, H. J. C.; Fraaije, J. G. E. M. LINCS: a linear constraint solver for molecular simulations. *J. Comput. Chem.* **1997**, *18*, 1463–1472.

(5) Berendsen, H. J. C.; Postma, J. P. M.; van Gunsteren, W. F.; DiNola, A.; Haak, J. R. Molecular dynamics with coupling to an external bath. *J. Chem. Phys.* **1984**, *81*, 3684–3690.

(6) Bussi, G.; Donadio, D.; Parrinello, M. Canonical sampling through velocity rescaling. *J. Chem. Phys.* **2007**, *126*, 014101.

(7) Parrinello, M.; Rahman, A. Polymorphic transitions in single crystals: A new molecular dynamics method. *J. Appl. Phys.* **1981**, *52*, 7182–7190.

(8) Bennett, C. H. Efficient estimation of free energy differences from Monte Carlo data. *J. Comput. Phys.* **1976**, *22*, 245–268.

(9) Hummer, G.; Pratt, L. R.; García, A. E. Free Energy of Ionic Hydration. *J. Phys. Chem.* **1996**, *100*, 1206–1215.

(10) Darden, T.; Pearlman, D.; Pedersen, L. G. Ionic charging free energies: Spherical versus periodic boundary conditions. *J. Chem. Phys.* **1998**, *109*, 10921–10935.

(11) Izadi, S.; Anandakrishnan, R.; Onufriev, A. V. Building water models: A different approach. *J. Phys. Chem. Lett.* **2014**, *5*, 3863–3871.

(12) Horinek, D.; Mamatkulov, S. I.; Netz, R. R. Rational design of ion force fields based on thermodynamic solvation properties. *J. Chem. Phys.* **2009**, *130*, 124507.

(13) Rick, S. W.; Stuart, S. J.; Berne, B. J. Dynamical fluctuating charge force fields: Application to liquid water. *J. Chem. Phys.* **1994**, *101*, 6141–6156.

(14) Lee Warren, G.; Patel, S. Hydration free energies of monovalent ions in transferable intermolecular potential four point fluctuating charge water: An assessment of simulation methodology and force field performance and transferability. *J. Chem. Phys.* **2007**, *127*.

(15) Beck, T. L. The influence of water interfacial potentials on ion hydration in bulk water and near interfaces. *Chem. Phys. Lett.* **2013**, *561-562*, 1–13.

(16) Tissandier, M. D.; Cowen, K. A.; Feng, W. Y.; Gundlach, E.; Cohen, M. H.; Earhart, A. D.; Coe, J. V.; Tuttle, T. R. The proton's absolute aqueous enthalpy and Gibbs free energy of solvation from cluster-ion solvation data. *J. Phys. Chem. A* **1998**, *102*, 7787–7794.

(17) Marcus, Y. *Ion Properties*; Marcel Dekker, Inc.: New York, Basel, 1997.

(18) Marcus, Y. Ionic Radii in Aqueous Solutions. *Chem. Rev.* **1988**, *88*, 1475–1498.

(19) Sengupta, A.; Li, Z.; Song, L. F.; Li, P.; Merz, K. M. Parameterization of Monovalent Ions for the OPC3, OPC, TIP3P-FB, and TIP4P-FB Water Models. *J. Chem. Inf. Model* **2021**, *61*, 869–880.

(20) Wynne-Jones, W. F. K.; Eyring, H. The Absolute Rate of Reactions in Condensed Phases. *J. Chem. Phys.* **1935**, *3*, 492–502.

(21) Wigner, E. The Transition State Method. *Trans. Faraday Soc.* **1937**, 29–41.

(22) Schwierz, N. Kinetic pathways of water exchange in the first hydration shell of magnesium. *J. Chem. Phys.* **2020**, *152*, 224106.

(23) Neely, J.; Connick, R. Rate of Water Exchange from Hydrated Magnesium Ion. *J. Am. Chem. Soc.* **1970**, *92*, 3476–3478.

(24) Bleuzen, A.; Pittet, P.-A.; Helm, L.; Merbach, A. E. Water exchange on magnesium(II) in aqueous solution: a variable temperature and pressure $^{17}$O NMR study. *Magn. Reson. Chem.* **1997**, *35*, 765–773.

(25) Torrie, G. M.; Valleau, J. P. Monte Carlo free energy estimates using non-Boltzmann sampling: Application to the sub-critical Lennard-Jones fluid. *Chem. Phys. Lett.* **1974**, *28*, 578–581.

(26) Torrie, G. M.; Valleau, J. P. Nonphysical sampling distributions in Monte Carlo free-energy estimation: Umbrella sampling. *J. Comput. Phys.* **1977**, *23*, 187–199.

(27) Abraham, M. J.; Murtola, T.; Schulz, R.; Páll, S.; Smith, J. C.; Hess, B.; Lindah, E. Gromacs: High performance molecular simulations through multi-level parallelism from laptops to supercomputers. *SoftwareX* **2015**, *1-2*, 19–25.

(28) Case, D. A.; Belfon, K.; Ben-Shalom, I. Y.; Brozell, S. R.; Cerutti, D. S.; Cheatham, T. E. I.; Cruzeiro, V. W. D.; Darden, T.; Duke, R. E.; Giambasu, G.; Gilson, M. K.; Gohlke, H.; Goetz, A. W.; Harris, R.; Izadi, P. A.; Izmailov, S.; Kasavajhala, K.; Kovalenko, A.; Krasny, R.; Kurtzman, T.; Lee, T. S.; LeGrand, S.; Li, P.; Lin, C.; Liu, J.; Luchko, T.; Luo, R.; Man, V.; Merz, K. M.; Miao, Y.; Mikhailovskii, O.; Monard, G.; Nguyen, H.; Onufriev, A.; Pan, F.; Pantano, S.; Qi, R.; Roe, D. R.; Roitberg, A.; Sagui, C.; Schott-Verdugo, S.; Shen, J.; Simmerling, C. L.; Skrynnikov, N. R.; Smith, J.; Swails, J.; Walker, R. C.; Wang, J.; Wilson, L.; Wolf, R. M.; Wu, X.; Xiong, Y.; Xue, Y.; York, D. M.; Kollman, P. A. Amber 2018. 2018; `https://ambermd.org/AmberTools.php`.

(29) Tribello, G. A.; Bonomi, M.; Branduardi, D.; Camilloni, C.; Bussi, G. Plumed 2: New feathers for an old bird. *Comput. Phys. Commun.* *185*, 604–613.

(30) Kumar, S.; Rosenberg, J. M.; Bouzida, D.; Swendsen, R. H.; Kollman, P. A. The weighted histogram analysis method for free-energy calculations on biomolecules. I. The method. *J. Comput. Chem.* **1992**, *13*, 1011–1021.

(31) Grotz, K. K.; Cruz-León, S.; Schwierz, N. Optimized Magnesium Force Field Parameters for Biomolecular Simulations with Accurate Solvation, Ion-Binding, and Water-Exchange Properties. *J. Chem. Theory Comput.* **2021**, *17*, 2530–2540.

(32) Grotz, K. K.; Schwierz, N. Optimized Magnesium Force Field Parameters for Biomolecular Simulations with Accurate Solvation, Ion-Binding, and Water-Exchange Properties in SPC/E, TIP3P-fb, TIP4P/2005, TIP4P-Ew, and TIP4P-D. *J. Chem. Theory Comput.* **2022**, *19*, 526–537.

(33) Kirkwood, J. G.; Buff, F. P. The Statistical Mechanical Theory of Solutions. I. *J. Chem. Phys.* **1951**, *19*, 774–777.

(34) Fyta, M.; Netz, R. R. Ionic force field optimization based on single-ion and ion-pair solvation properties: Going beyond standard mixing rules. *J. Chem. Phys.* **2012**, *136*, 124103.

(35) Weerasinghe, S.; Smith, P. E. A Kirkwood-Buff derived force field for sodium chloride in water. *J. Chem. Phys.* **2003**, *119*, 11342–11349.

(36) Sigel, R. K.; Sigel, H. A stability concept for metal ion coordination to single-stranded nucleic acids and affinities of individual sites. *Acc. Chem. Res.* **2010**, *43*, 974–984.

(37) Leonarski, F.; D'Ascenzo, L.; Auffinger, P. $Mg^{2+}$ ions: Do they bind to nucleobase nitrogens? *Nucleic Acids Res.* **2017**, *45*, 987–1004.

(38) Chipot, C.; Pohorille, A. *Free energy calculations*; Springer Science Business Media: Heidelberg, Berlin, 2007.

(39) Cruz-León, S.; Vanderlinden, W.; Müller, P.; Forster, T.; Staudt, G.; Lin, Y.; Lipfert, J.; Schwierz, N. Twisting DNA by Salt. *bioRxiv2021.07.14.452306* **2021**, 21–26.

(40) Shirts, M. R.; Chodera, J. D. Statistically optimal analysis of samples from multiple equilibrium states. *J. Chem. Phys.* **2008**, *129*, 1–10.

(41) Mamatkulov, S.; Schwierz, N. Force fields for monovalent and divalent metal cations in TIP3P water based on thermodynamic and kinetic properties. *J. Chem. Phys.* **2018**, *148*, 74504.
